# Ribosomal RNA transcription governs splicing through ribosomal protein RPL22

**DOI:** 10.1101/2024.08.15.608201

**Authors:** Wenjun Fan, Hester Liu, Gregory C. Stachelek, Asma Begum, Catherine E. Davis, Tony E. Dorado, Glen Ernst, William C. Reinhold, Busra Ozbek, Qizhi Zheng, Angelo M. De Marzo, N.V. Rajeshkumar, James C. Barrow, Marikki Laiho

## Abstract

Ribosome biosynthesis is a cancer vulnerability executed by targeting RNA polymerase I (Pol I) transcription. We developed advanced, specific Pol I inhibitors to identify drivers of this sensitivity. By integrating multi-omics features and drug sensitivity data from a large cancer cell panel, we discovered that *RPL22* frameshift mutation conferred Pol I inhibitor sensitivity in microsatellite instable cancers. Mechanistically, RPL22 directly interacts with 28S rRNA and mRNA splice junctions, functioning as a splicing regulator. RPL22 deficiency, intensified by 28S rRNA sequestration, promoted the splicing of its paralog RPL22L1 and p53 negative regulator MDM4. Chemical and genetic inhibition of rRNA synthesis broadly remodeled mRNA splicing controlling hundreds of targets. Strikingly, RPL22-dependent alternative splicing was reversed by Pol I inhibition revealing a ribotoxic stress-initiated tumor suppressive pathway. We identify a mechanism that robustly connects rRNA synthesis activity to splicing and reveals their coordination by ribosomal protein RPL22.

## Introduction

Ribosome biogenesis is indispensable for cancer cell growth (Grummt 2010; Ruggero 2012; Pelletier et al. 2018). The first and rate-limiting step in ribosome biosynthesis is RNA polymerase I (Pol I) transcription, and in most cancers the transcription is pervasively activated. Key oncogenic drivers such as MYC, RAS/ERK, AKT/PKB, and mTOR upregulate Pol I transcription and conversely, the transcription is repressed by tumor suppressors most frequently inactivated in cancers, namely p53, RB1, CDKN2A (ARF) and PTEN (Grummt 2010; Ruggero 2012; Bywater et al. 2013; Pitts and Laiho 2022). Despite this knowledge, Pol I has remained an underexplored target for selective inhibition of cancer cell growth. However, several chemotherapeutic drugs repress Pol I transcription (Burger et al. 2010; Bywater et al. 2013; Pitts and Laiho 2022; Zisi et al. 2022). Most inhibit Pol I indirectly by targeting topoisomerases essential for resolving torsional stress of the heavily transcribed rRNA genes (Burger et al. 2010; Bruno et al. 2017; Pitts and Laiho 2022). This shortcoming hampers the understanding of the contribution of topoisomerase inhibition versus Pol I inhibition towards anticancer efficacy and compromises the identification of sensitivity drivers. CX-5461, previously nominated as specific Pol I inhibitor (Bruno et al. 2017), has unequivocally been identified as a TOP2 targeting agent and conveys its therapeutic activity by this mechanism (Bruno et al. 2020; Bossaert et al. 2021; Pan et al. 2021; Xu and Hurley 2022).

We have identified structurally distinct Pol I-targeting molecules that block three critical steps in Pol I transcription cycle, namely initiation, promoter escape, and elongation (Colis et al. 2014a; Peltonen et al. 2014a; Peltonen et al. 2014b; Wei et al. 2018; Dorado et al. 2022; Jacobs et al. 2022a; Jacobs et al. 2022b). Consequently, the inhibitors activate proteasome-dependent degradation of the enzyme catalytic subunit by SCF^FBXL14^ E3 ligase (Peltonen et al. 2014b; Pitts et al. 2022). Pol I degradation is only observed in transcription-competent cells, linking polymerase stability to transcriptional activity (Wei et al. 2018). The activity of the inhibitors is genetically dependent on the Pol I enzyme complex in cancer cells and in yeast, confirming Pol I as their direct and specific target (Wei et al. 2018; Low et al. 2019; Ford et al. 2023). Furthermore, genome-wide and *in vitro* analyses show that the inhibitors are selective against Pol I compared to Pol II and Pol III (Jacobs et al. 2022a; Jacobs et al. 2022b). Lastly, in contrast to the chemotherapeutic Pol I inhibitors, these direct Pol I inhibitors do not cause DNA damage (Peltonen et al. 2010; Colis et al. 2014a; Colis et al. 2014b; Peltonen et al. 2014a; Peltonen et al. 2014b; Xu et al. 2017; Bruno et al. 2020).

Inhibition of Pol I transcription activates a ribotoxic stress pathway that culminates in the activation of p53 (Deisenroth et al. 2016; Pelletier et al. 2018). This pathway is orchestrated by ribosomal proteins RPL5 and RPL11 that block the ability of MDM2 to ubiquitinate p53 (Lohrum et al. 2003; Bursać et al. 2012; Deisenroth et al. 2016; Pelletier et al. 2018). However, this knowledge has not been therapeutically exploitable given the lack of alterations of RPL5 and RPL11 in cancers. Here, we report the discovery of an alternative ribotoxic stress mechanism that operates parallel to the RPL5/RPL11 regulatory circuitry. Using cancer cell line screens and by conducting correlative analyses for molecular features associated with Pol I inhibitor sensitivity, we identified an inactivating mutation in a tumor suppressor and ribosomal protein *RPL22*, and high expression of its paralog RPL22L1 and p53 repressor MDM4 as markers for sensitivity to the inhibitors. These markers were enriched in microsatellite instable (MSI) cancer cell lines, and cell lines and tumor models bearing these markers were highly responsive to Pol I inhibition. We demonstrate that RPL22 binds hundreds of mRNAs and alters their splicing. We find that Pol I inhibition regulates the splicing of a multitude of mRNAs and specifically, RPL22L1 and MDM4 and that this activity is conveyed by RPL22. Our data reveal that high rRNA synthesis activity sequesters RPL22 into the ribosomes and counters the RPL22-mediated splicing regulation. Thus, RPL22 coordinates rRNA synthesis with splicing programs. These findings provide a molecular mechanism that connects rRNA synthesis activity with post-transcriptional and spliceosomal control and reveal the perturbations of this axis in human cancers.

## Results

### Cancer cell line screens for Pol I inhibition identify novel ribotoxic stress response factors

We assessed the growth inhibition 50 (GI_50_) of the tool compound BMH-21 and an optimized derivative, BOB-42 (Colis et al. 2014a), in a panel of over 300 cancer cell lines representing 18 lineages (**Figure 1A**). The responses of cell lines to BOB-42 and BMH-21 were highly similar with Pearson correlation (*r*) over 0.9 (**Figure 1B**). The most sensitive lineages were leukemia, kidney, colorectal, myeloma, pancreas and ovarian, but exceptional responders were observed in all (**Figure 1C**; **Tables S1** and **S2**). Cell lines that were MSI-high due to defects in mismatch repair (dMMR), were significantly more sensitive to the inhibitors than microsatellite stable (MSS) lines (**Figure 1D**).

**Figure 1.**
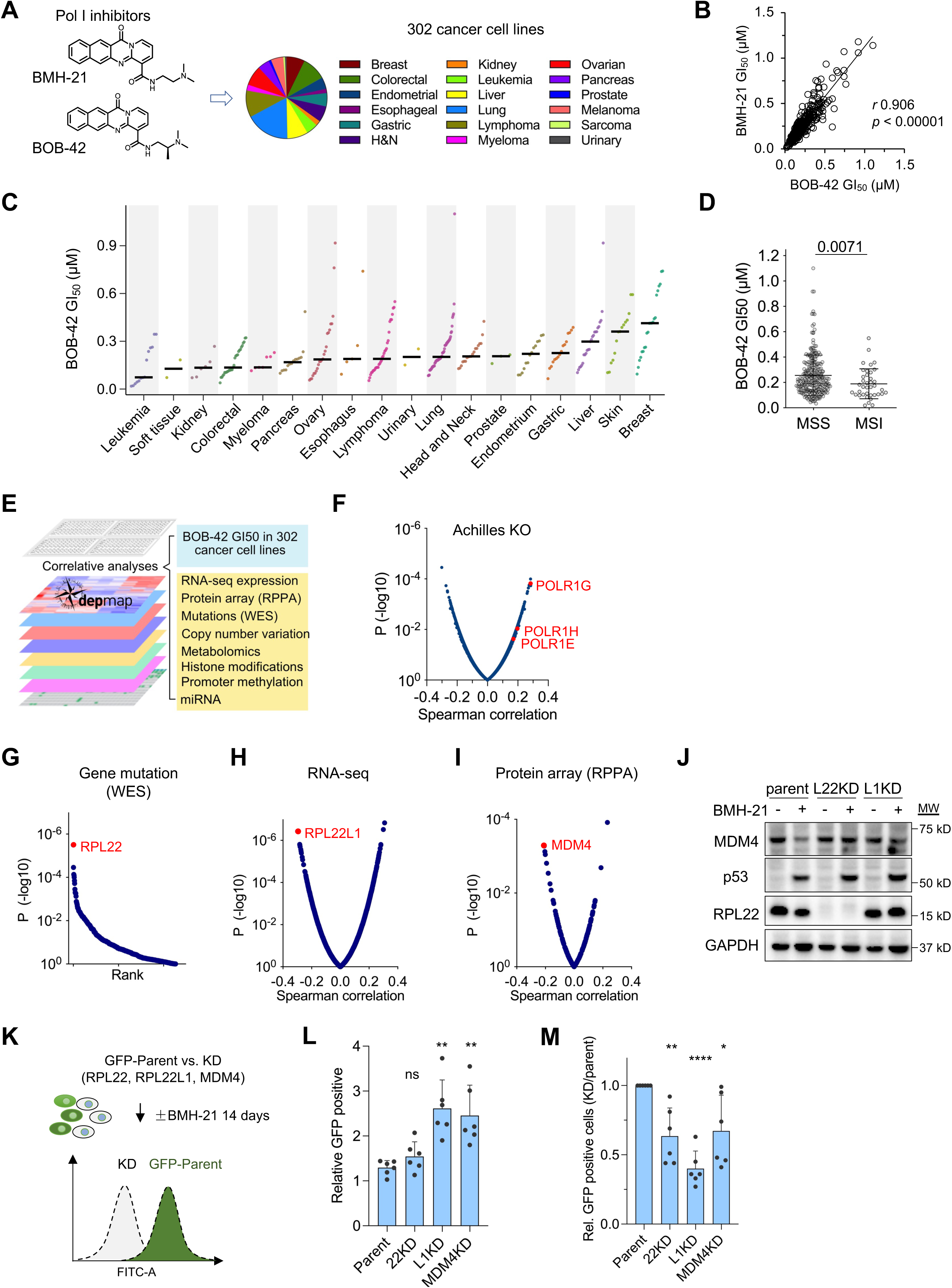
Cancer cell line screens identify markers for sensitivity to Pol I inhibitors. (**A**) Schematic of cancer cell line screens. (**B**) Pearson correlation analyses for BMH-21 and BOB-42 growth inhibitory dose 50 (GI_50_) for (n = 302 cell lines). (**C**) CellTiterGlo viability assay for BOB-42. GI_50_ (µM) is shown. **D**, Dependency of BOB-42 GI_50_ on microsatellite stable (MSS) and instable (MSI) status (n = 291 cell lines). *p*, two-tailed Mann-Whitney test. (**E**) Schematic of correlative analyses of BOB-42 cancer cell line response profiles with DepMap features. (**F**) BOB-42 GI_50_ efficacy was correlated to Achilles database. Pol I subunit genes are shown (n = 184 cell lines). (**G**) Gene mutations ranked based on significance (whole exome sequencing, WES). (**H and I**) Spearman correlations of BOB-42 GI_50_ with (**H**) gene expression (RNA-seq) (n = 280 cell lines), and (**I**) protein expression (RPPA) (n = 270 cell lines). (**J**) Immunoblotting of HCT116 parent, RPL22 (L22KD) and RPL22L1 (L1KD) knockdown cells. Representative experiment of n = 3 is shown. (**K**) Schematic outline of GFP-competition growth assay. (**L and M**) Competition assay of GFP-labelled parent cells mixed with unlabeled MDM4^KD^, RPL22^KD^, and RPL22L1^KD^ cells followed by flow cytometry. **L**, The fraction of GFP-positive parent cells is shown. (**M**) Cells were treated with BMH-21 (0.25 µM) for 14 days. The relative fold change of GFP-positive cells was normalized to the parent cells set as 1. n = 6 independent biological replicates, mean±SD. *p* values, Student’s two-sided t-test. *ns*, non-significant, * <0.05, ** <0.01, **** <0.0001. See also Figure S1, Tables S1 and S2.

Molecular features associated with the cancer cell line responses were interrogated by correlative analyses using DepMap database (**Figure 1E**) (Tsherniak et al. 2017). Features associated with Pol I inhibitor sensitivity showed several strong and statistically highly significant correlations. Comparison to the Achilles gene essentiality database showed high correlation with CRISPR-Cas9 knockout (KO) of three Pol I complex subunits (**Figure 1F**). *RPL22^K15Rfs^* mutation was the top-ranking mutational event (**Figure 1G**) and *RPL22* mutant lines were significantly more sensitive to BOB-42 (**Figure S1A**). These were all frameshift mutations affecting almost exclusively *RPL22^K15R^*, which is a hotspot mutation present in up to 70% MSI cancer cell lines and dMMR tumors (Ferreira et al. 2014; Maruvka et al. 2017; Ghandi et al. 2019). Interrogation of the RNA-seq database for transcript expression identified RPL22L1 as the top-ranking transcript correlating with sensitivity (**Figure 1H**). This finding is robust, as we observed a similar correlation with BMH-21 response in an independent NCI60 cancer cell line dataset (**Figure S1B**). RPL22 and RPL22L1 are a synthetic lethal paralog pair. RPL22 is a negative regulator of RPL22L1 and consequently, haploinsufficiency or loss of *RPL22* increases RPL22L1 (O’Leary et al. 2013; McDonald et al. 2017; Rao et al. 2019). Accordingly, we found that lines with *RPL22* mutation had significantly higher expression of RPL22L1 (**Figure S1C**). Finally, the top-ranking protein correlate was MDM4 (**Figure 1I**). Given the sensitivity of MSI-high cancer cell lines to the inhibitors, we compared the expression of MDM4 protein in MSS and MSI lines. As shown in **Figure S1D**, MDM4 protein was significantly higher in MSI lines. Also, the expression of MDM4 full-length (FL) splice variant and wild-type *TP53* were more frequent in the MSI lines compared to MSS lines (**Figures S1E** and **S1F**).

In contrast, neither *TP53* mutation (*p* 0.252) nor the expression of RPL5 (*p* 0.290) or RPL11 (*p* 0.0513) reached statistical significance in these analyses. This is consistent with the observation that while *TP53* wild-type cancer cell lines responded well to the Pol I inhibitors, p53 is neither required nor the sole driver of sensitivity to BMH-21 (Peltonen et al. 2014b). We then tested whether RPL22 and RPL22L1 function in the classical ribotoxic stress pathway and stabilize p53. Strikingly, their depletion did not alter the abundance of p53, although p53 was robustly stabilized following treatment with the Pol I inhibitor (**Figure 1J**). This shows that neither RPL22 nor RPL22L1 function in the known ribotoxic stress response pathway stabilizing p53 through MDM2. This is consistent with observations of their roles in mouse developmental models (Solanki et al. 2016; Fahl et al. 2022). These findings identify that RPL22, RPL22L1 and MDM4 correlate with sensitivity to Pol I inhibition and plausibly converge to a distinct ribotoxic stress pathway.

Next, we used a GFP-competition assay to examine the marker-dependent sensitivity to BMH-21. We used HCT116 MSI colorectal cancer cell line, which harbors the *RPL22^K15Rfs^* mutation, and generated stable knock-down lines for each of RPL22, RPL22L1 and MDM4 (**Figures S1G–S1I**). GFP-labelled parent HCT116 cells were mixed at 1:1 ratio with either unlabeled parent, RPL22^KD^, RPL22L1^KD^ or MDM4^KD^ cells followed by treatment with BMH-21 and analysis by flow cytometry (**Figure 1K**). RPL22L1^KD^ and MDM4^KD^ caused a significant growth delay, suggesting that they promote cancer cell growth (**Figure 1L**). The knockdown of RPL22, MDM4 and RPL22L1 each led to significant resistance to Pol I inhibition (**Figure 1M**).

### Pol I inhibitor is highly effective in MDM4-high and *RPL22*-mutant preclinical mouse models

Our preclinical studies *in vivo* have shown effective targeting of cancers such as melanoma, colon and prostate by BMH-21 (Low et al. 2019; Fahl et al. 2022). BOB-42 is an advanced molecule with optimal drug-like properties and pharmacokinetics suitable for preclinical studies (**Figure S2A**) (Colis et al. 2014a). We conducted *in vivo* efficacy studies in models expressing the identified markers. First, BOB-42 dose-dependent efficacy was tested in a MDM4-high A375 melanoma xenograft model. As shown in **Figure 2A**, BOB-42 effectively inhibited tumor growth by up to 77%. The treatment was well tolerated without mouse weight loss or alterations in clinical chemistry (**Figure S2B** and **Table S3**). Complete blood count differential analysis showed that while there was no change in white blood cells or platelets, the treatment caused a dose-dependent decrease in reticulocytes (**Table S4**). This was not considered rate-limiting for therapeutics. A further dose-response study using alternative dosing schedules showed the highest efficacy (tumor growth inhibition 90%) at BOB-42 100 mg/kg (**Figure S2C**). BOB-42 tumor uptake was excellent (30-100-fold as compared to plasma levels) and strongly correlated with tumor growth control (**Figure S2D**). Furthermore, the exposure strikingly correlated with a reduction in Pol I transcription demonstrating target engagement, as shown by quantitative *in situ* hybridization for 45S precursor rRNA (**Figure S2E**).

**Figure 2.**
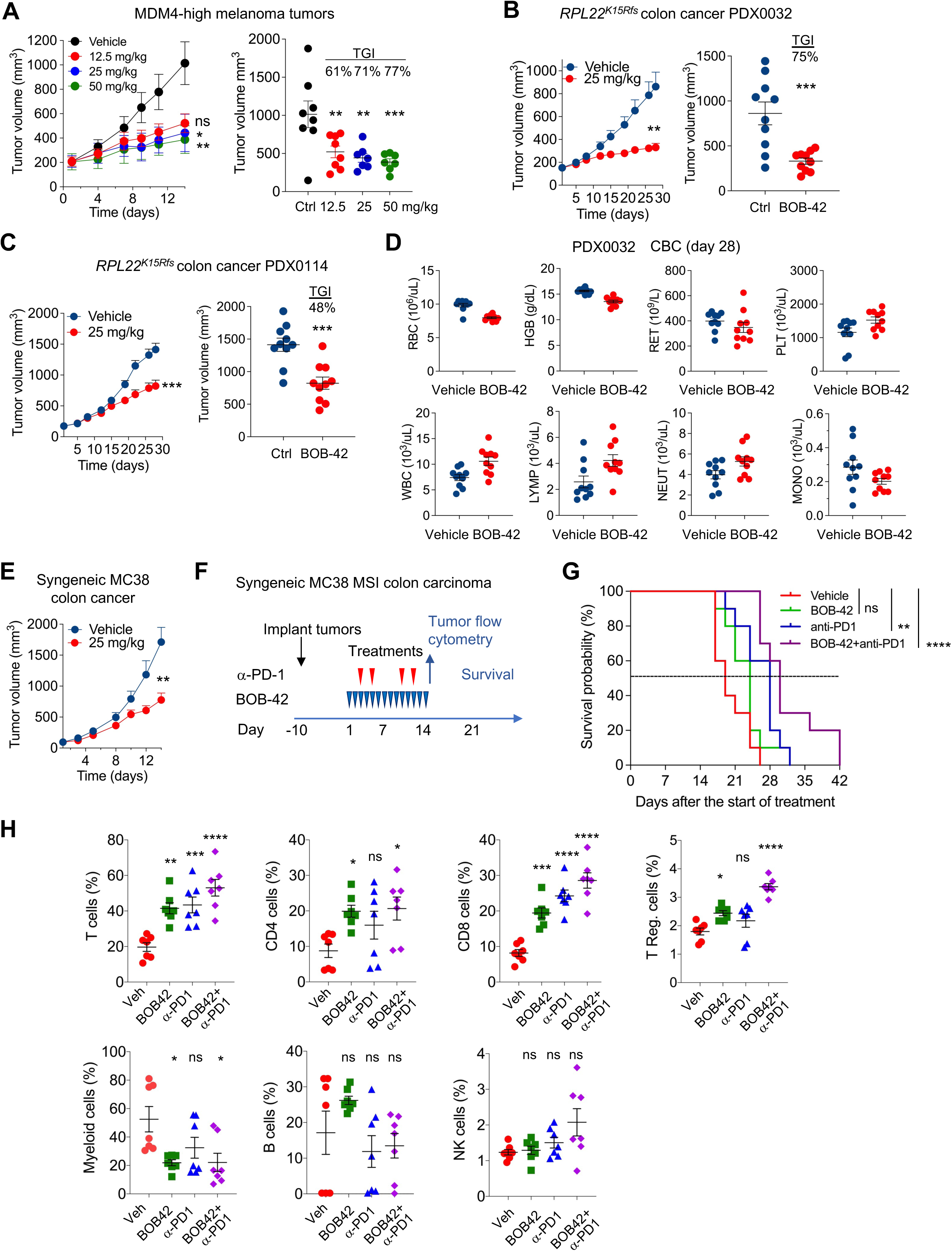
BOB-42 *in vivo* efficacy in tumor models expressing the sensitivity markers. (**A**) Tumor growth curves and volumes of MDM4-high A375 melanoma xenografts following treatments with BOB-42. n = 8 mice/group. TGI, tumor growth inhibition. Whiskers and bars represent mean±SEM. *p* values, One-Way ANOVA with Tukey’s multiple comparison test. (**B and C**) Tumor growth curves and volumes of human *RPL22^K15Rfs^*MSI-high colon carcinoma patient-derived xenografts (PDX) following treatment with BOB-42 (25 mg/kg). **B**, PDX0032. **C**, PDX0114. n = 10 mice/group. Whiskers and bars represent mean±SEM. *p* values, unpaired two-sided t-test with Welch’s correction. (**D**) Complete blood counts in (B). Mean±SEM. (**E**) Tumor growth curves of syngeneic MC38 xenografts following treatment with BOB-42 (25 mg/kg). n = 10 mice/group. *p* values, One-way ANOVA with Dunnett’s two-sided analysis. (**F**) Schematic diagram of the study. BOB-42 (*blue* triangles) and anti-PD1 antibody (*red* triangles). (**G**) Kaplan-Mayer survival curves. n = 10 mice/group. *p* values, Log-rank (Mantel-Cox) test. (**H**) Flow cytometry analysis of immune cells. The frequency of CD45^+^ cells are shown. n = 7 mice/group. *p* values, Tukey’s multiple comparison test. *p* values, *ns*, non-significant, * <0.05, ** <0.01, *** <0.001, **** <0.0001. See also Figure S2, Tables S3 and S4.

We then tested the efficacy of BOB-42 in two human MSI colon carcinoma patient-derived xenografts (PDX) with *RPL22^K15Rfs^* mutation, which showed up to 75% tumor growth control (**Figures 2B** and **2C**). No toxicity signals were detected. There was no effect of the daily 4-week BOB-42 treatment on mouse weight (**Figures S2F** and **S2G**), well-being, or complete blood count (**Figure 2D**).

We further tested the effect of BOB-42 in MC38 syngeneic colorectal cancer model in immunocompetent *C57BL/6* mice. MC38 tumors carry mutations in *Tp53*, mismatch repair gene *Msh3* and DNA replication/ repair gene *Pold1,* have high mutation burden, and are considered mismatch repair defective^39^. As shown in **Figure 2E**, BOB-42 reduced the growth of MC38 syngeneic tumors by 50%, while there were no adverse effects on mouse weight (**Figure S2H**). Given the clinically successful use of immune checkpoint inhibitors (ICI) in dMMR cancers (Colis et al. 2014a), we assessed the contribution of immune responses to the efficacy of BOB-42 with and without ICI. We treated established MC38 tumors with BOB-42 and α-PD-1 antibodies using the treatment scheme in **Figure 2F** and measured in-tumor immune cell responses and mice survival. BOB-42 alone did not increase survival, and while α-PD-1 modestly did so, the combination had a highly significant survival advantage (**Figure 2G**). Flow cytometry analyses showed increased intratumor frequency of T cells and CD8^+^/CD4^+^ T cells by BOB-42 alone, which was further augmented by the ICI combination (**Figure 2H**). Remarkably, BOB-42 alone and in combination with α-PD-1 decreased myeloid cell infiltration (**Figure 2H**).

### Pol I inhibition decreases the expression of RPL22L1 and regulates its splicing

High expression of RPL22L1 correlated with drug sensitivity (**Figure 1H**). We hypothesized that the expression of RPL22L1 is regulated by Pol I inhibition. Testing this in a kinetic experiment showed that RPL22L1 was repressed by 80% by BMH-21 at 16 hours (**Figure 3A**). We then treated *TP53* proficient (p53^+/+^) and deficient (p53^-/-^) HCT116 isogenic cells with BMH-21 and found that the treatment profoundly decreased RPL22L1 expression in both lines (**Figure 3B**). However, BMH-21 caused only a minor 20% decrease of RPL22 mRNA and protein in the parent cells (**Figures 3C–3E**). Given that RPL22 and RPL22L1 are a synthetic lethal paralog pair (McDonald et al. 2017), we also tested whether the expression of RPL22 is under the control of RPL22L1. RPL22L1 was effectively knocked out using sgRNAs (**Figure S1J**). RPL22 expression and protein abundance increased by 2-fold in the RPL22L1^KO^ cells, and this increase was unaffected by BMH-21 (**Figure 3C–E**). To further assess the regulation of RPL22L1 by Pol I inhibition, we analyzed its expression in a panel of colon, stomach, prostate and endometrial cancer cell lines that were either wild type for *RPL22* (3 lines) or had *RPL22^K15Rfs^* mutation (7 lines). BMH-21 significantly decreased RPL22L1 in each line, irrespective of *RPL22* or *TP53* mutation (**Figure 3F**).

**Figure 3.**
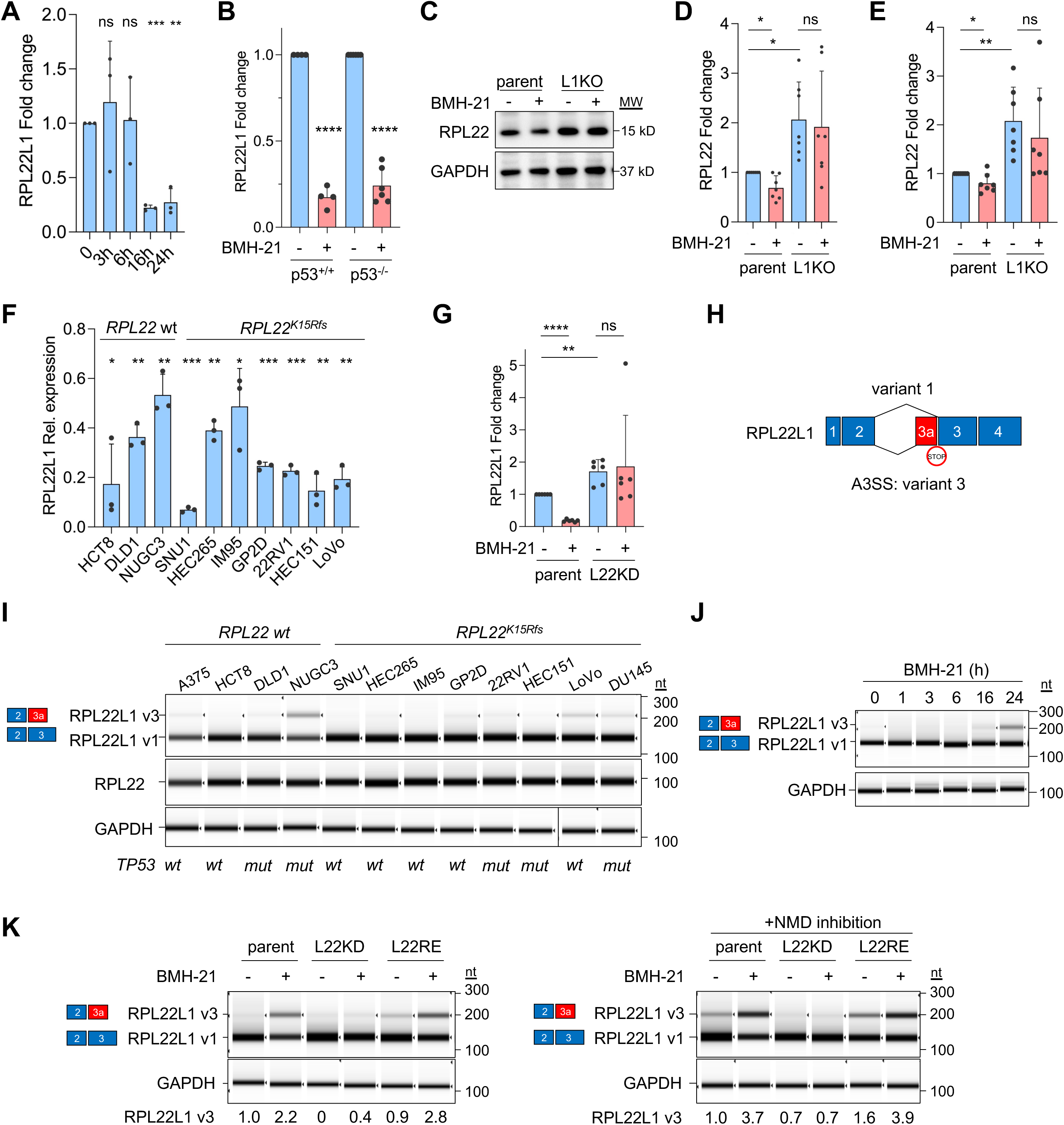
Pol I inhibition decreases the expression of RPL22L1. (**A**) qPCR for RPL22L1 in HCT116 p53^+/+^ cells treated with BMH-21 (1 µM) for the indicated times. (**B**) qPCR for RPL22L1 in HCT116 *TP53* isogenic cells treated with BMH-21 (1 µM). (**C and D**) Immunoblotting and quantification of RPL22. (**E**), qPCR for RPL22 in parent and RPL22L1^KO^ cells. (**F**) qPCR for RPL22L1 in cancer cell lines wild type (wt) or mutant for *RPL22*. *TP53* mutant cell lines are indicated. (**G**) qPCR for RPL22L1 in parent and RPL22^KD^ cells. (**H**) Schematic of RPL22L1 splicing. (**I**) RPL22L1 isoforms in cancer cell lines. *RPL22* genotypes and *TP53* mutant lines are indicated. Representative sqPCR is shown. (**J**) RPL22L1 isoforms in p53^+/+^ cells. Representative sqPCR of n = 2 is shown. (**K**) RPL22L1 isoforms in RPL22^KD^ and RPL22^RE^ cells treated with BMH-21 and NMD inhibitor anisomycin. Representative sqPCR of n = 2 is shown. Data are representative of at least three independent experiments (A - G) and mean±SD are shown. *P* values, Student’s two-sided t-test. *ns*, non-significant, * <0.05, ** <0.01, *** <0.001, **** <0.0001.

RPL22 negatively regulates RPL22L1 (O’Leary et al. 2013). We hence tested whether the regulation of RPL22L1 by Pol I inhibition depends on RPL22. This question was not fully answerable in the *RPL22^K15Rfs^* mutant lines, as these are heterozygous for *RPL22* (see **Figure 3I**). We used stable RPL22 knockdown (RPL22^KD^) cells to assess this dependency. RPL22^KD^ led to a moderate but significant compensatory increase in RPL22L1 (**Figure 3G**). Remarkably, the decrease in RPL22L1 by BMH-21 was abrogated by RPL22 knockdown (**Figure 3G**).

RPL22L1 undergoes alternative splicing (Larionova et al. 2022; Weinstein et al. 2023). Splicing of an alternative 3’ splice site in intron 2 leads to a frameshift and expression of a putative 64 amino acid product with an early stop codon that lacks the RPL22L1 RNA interaction domain and is termed as variant 3 here (**Figure 3H**). Variant 3 has been detected in glioblastoma cells (Larionova et al. 2022). We analyzed the main isoform (called variant 1) and variant 3 using splicing specific primers by semiquantitative PCR (sqPCR) in cancer cell lines and identified constitutive, albeit low expression of variant 3 in several lines independent of *RPL22* mutation (**Figure 3I**). We then analyzed RPL22L1 isoform regulation by Pol I inhibition and observed increased variant 3 expression following 16-hour BMH-21 treatment (**Figure 3J**). We then asked whether the splicing is dependent on RPL22. RPL22 knockdown led to the complete abrogation of RPL22L1 splicing by Pol I inhibition, whereas the reconstitution of RPL22 in the *RPL22^K15Rfs^* mutant cells increased the expression of variant 3 by BMH-21 (**Figure 3K**). Variant 3 expression was further stabilized by anisomycin, which inhibits nonsense-mediated decay (NMD) (**Figure 3K**) (Hauser et al. 2020). Together, these findings demonstrated that RPL22L1 variant 1 and 3 expression is oppositely regulated by Pol I inhibition in a RPL22-dependent manner and showed that Pol I inhibition predominantly affects RPL22L1 but not RPL22 abundance.

### Repression of rRNA synthesis decreases MDM4 by altering its splicing

Ribotoxic stress stabilizes p53 by inhibition of MDM2 by RPL5 and RPL11 (Pelletier et al. 2018). Previous studies showed that MDM4 protein stability was decreased by ribotoxic stress (Gilkes et al. 2006; Li and Gu 2011). We first analyzed changes in MDM4 protein abundance by using BMH-21. MDM4 protein robustly decreased and became undetectable by 16 h in both HCT116 p53 isogenic cell lines (**Figure 4A**). In parallel, p53 was stabilized and p53 target genes CDKN1A and MDM2 were induced in p53^+/+^ but not in p53^-/-^ cells indicating transcriptional activation of p53 (**Figure 4A**, **Figures S3A** and **S3B**).

**Figure 4.**
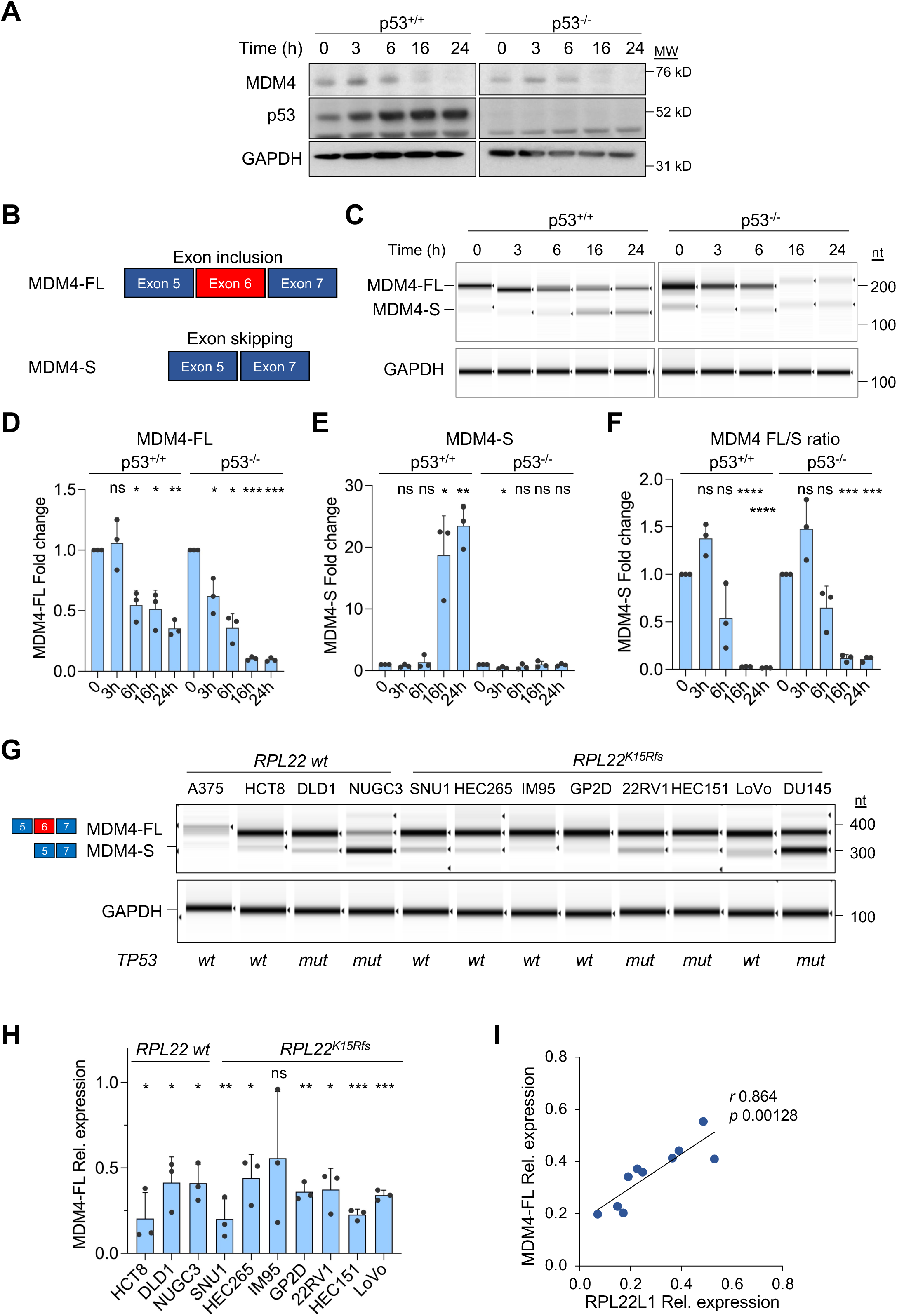
Pol I inhibitors alter MDM4 splicing and repress MDM4-FL. (**A**) Immunoblotting of HCT116 isogenic cells. (**B**) Schematic diagram of MDM4 splicing. (**C**) SqPCR analysis for MDM4-FL and MDM-S isoforms in p53 isogenic cells. (**D -F**) p53^+/+^ and p53^-/-^ cells were treated with BMH-21 (1 µM) for the indicated times. **D**, MDM4-FL and **E**, MDM-S isoforms were measured by qPCR and their ratios are shown in **F**. (**G**) MDM4 isoforms in cancer cell lines. Representative sqPCR of n = 2 is shown. (**H**) qPCR for MDM4-FL in cancer cell lines. *RPL22* genotypes and *TP53* mutant lines are indicated. (**I**) Correlation of regulation of MDM4-FL and RPL22L1 by BMH-21. *Inset*, Pearson *r* and statistical significance. Data are representative of at least three independent experiments (A, C, D -F, H) and mean±SD are shown. *p* values, Student’s two-sided t-test. *ns*, non-significant, * <0.05, ** <0.01, *** <0.001, **** <0.0001. See also Figure S3.

Earlier studies linked RPL22, RPL22L1 and MDM4 with sensitivity to MDM2 inhibitor Nutlin-3a (Ghandi et al. 2019; Weinstein et al. 2023). However, these relationships were not mechanistically investigated. Alternative splicing of MDM4 generates MDM4 full-length (FL) and MDM4 short (S) isoforms by inclusion or skipping of exon 6, respectively (**Figure 4B**) (Lenos et al. 2012; Bezzi et al. 2013; Dewaele et al. 2016; Marine and Jochemsen 2016). MDM4-FL represses p53, whereas MDM4-S transcript undergoes NMD (Dewaele et al. 2016). We hypothesized that Pol I stress could impact MDM4 splicing. We analyzed MDM4 splicing by using sqPCR and qPCR using splicing specific primers in p53^+/+^ and p53^-/-^ cells. As shown in **Figures 4C–4E**, BMH-21 robustly decreased the expression of MDM4-FL by 80% in both lines. MDM4-S transcript increased by over 20-fold, but only in p53 proficient cells (**Figure 4E**). This showed that MDM4-FL splicing was independent of p53, while that of MDM4-S was not. MDM4-FL/S ratio, which robustly predicts the expression of MDM4 protein,^48^ was profoundly repressed in both p53 isogenic lines (**Figure 4F**).

We then tested MDM4 splicing in ten cancer cell lines that were either *RPL22* wild-type or mutant. First, analysis of these lines by sqPCR showed that they highly expressed MDM4-FL, and that *TP53* mutant lines also highly expressed MDM4-S consistent with the earlier observation that *TP53* mutation was associated with MDM4 splicing change (**Figure 4G**) (Ghandi et al. 2019). Second, BMH- 21 decreased the expression of MDM4-FL in all lines irrespective of *RPL22* or *TP53* genotype (**Figure 4H**). However, the decrease in MDM4-FL strongly and significantly correlated with the decrease in RPL22L1 (Pearson *r* 0.864, *p* 0.00128) shown in **Figure 3F** highlighting their coordinate regulation by Pol I inhibition (**Figure 4I**).

To further explore the regulation of RPL22L1 and MDM4-FL we used a broad-spectrum Pol I inhibitor actinomycin D, PRMT5 splicing inhibitor GSK3326595, and CLK-kinase inhibitor TG003 that inhibits SRS-splicing factor phosphorylation (Muraki et al. 2004; Gerhart et al. 2018), and compared these to MDM2 inhibitor Nutlin-3a. Temporal analysis of these chemical perturbations showed that only actinomycin D reduced both RPL22L1 and MDM4-FL, whereas GSK3326595 and Nutlin-3a were largely without effect on either MDM4 isoform or RPL22L1 (**Figure S3C**). TG003 robustly increased the expression of MDM4-S consistent with its ability to inhibit MDM4-regulating splicing factors (**Figure S3C**) (Bezzi et al. 2013). These data underscore that Pol I inhibitors possess an exceptional capacity to inhibit both MDM4 and RPL22L1.

### MDM4 splicing is dependent on RPL22

Earlier studies in developmental models showed that RPL22 causes exon skipping and RPL22L1 exon inclusion of Smad2 (Zhang et al. 2017). Having ascertained that Pol I activity regulates MDM4 isoform expression, we asked whether RPL22 or RPL22L1 regulate MDM4 splicing. For unbiased assessment of splicing, we conducted RNA-seq followed by the analysis of splicing alterations using rMATS (Shen et al. 2014) in HCT116 parent cells, cells reconstituted with RPL22 expression (RPL22^RE^), and RPL22L1^KO^ cells (**Figure 5A** and **Figure S4A**). We uncovered substantial alterations in RNA transcript expression and broad changes in splicing in RPL22^RE^ and RPL22L1^KO^ cells (**Figures S4B** and **S4D**). Over 1,500 splicing events were detected in each line, and these were predominantly exon-skipping events (**Figures 5B** and **5C** and **Tables S5** and **S6**). Splicing of MDM4 was a highly significant event in both RPL22L1^KO^ and RPL22^RE^ cells, and Sashimi plot analysis showed an increase in exon 6 skipping (**Figures 5D – 5F**). We then used these cells to analyze basal and BMH-21- regulated splicing of MDM4. Analysis of the splicing variants using sqPCR showed that the basal expression of MDM4-FL was reduced, and the expression of MDM4-S increased in RPL22L1^KO^ cells (**Figure 5G**). The strong repression of MDM4-FL by BMH-21 was not rescued. However, RPL22 knockdown, but not its reconstitution, completely abrogated the BMH-21-mediated decrease of MDM4-FL (**Figure 5H**). These findings were corroborated by analyses of the MDM4 splice variants by using qPCR and showed that MDM4 splicing by BMH-21 depended on RPL22 but was independent on RPL22L1 (**Figures 5I** and **5J**). However, as RPL22 has no splicing function it was ambiguous how this dependency was conveyed.

**Figure 5.**
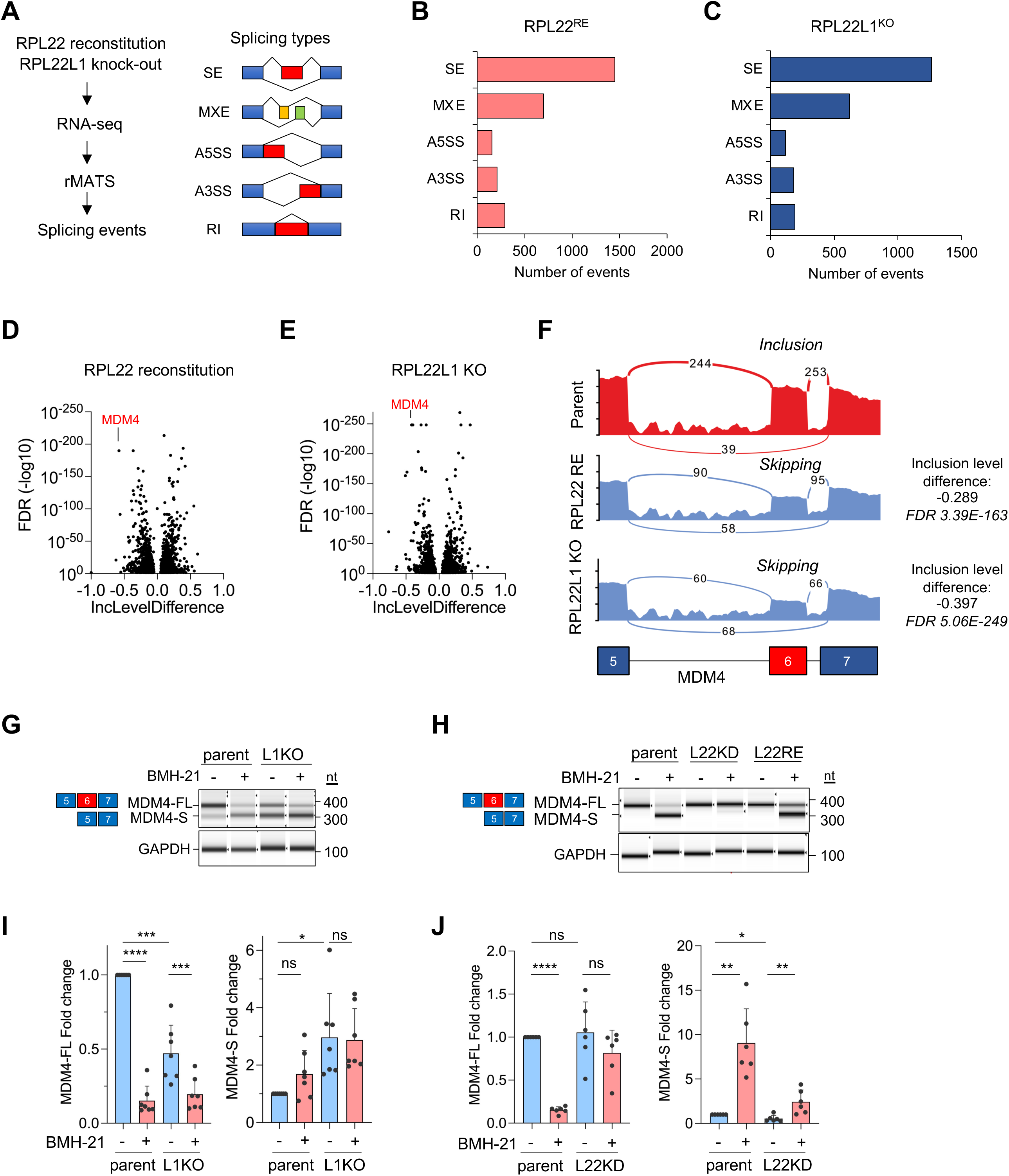
RPL22 is essential for regulation of MDM4 splicing by Pol I inhibition. (**A**) Schematic diagram of workflow and splicing events. SE, skipped exon; MXE, mutually exclusive exon; A5SS, alternative 5’ splice site; A3SS, alternative 3’ splice site; RI, retained intron. (**B**) Splicing events in RPL22^RE^ cells. (**C**) Splicing events in RPL22L1^KO^ cells. (**D and E**) Volcano plots of splicing events in RPL22^RE^ and RPL22L1^KO^ cells. (**F**) Sashimi plot of MDM4 splicing in RPL22^RE^ and RPL22L1^KO^ cells. (**G**) MDM4 isoforms in RPL22L1^KO^ cells. (**H**) MDM4 isoforms in RPL22^RE^ and RPL22^KD^ cells. A representative sqPCR of n = 2 biological replicates is shown. (**I**) qPCR for MDM4 isoforms in parent and RPL22L1^KO^ cells. (**J**) Same as in **I**, RPL22^KD^ cells. Data are representative of at least three independent experiments (G, I, J) and mean±SD are shown. *p* values, Student’s two-sided t-test. *ns*, non-significant, * <0.05, ** <0.01, *** <0.001, **** <0.0001. See also Figure S4, Tables S5 and S6.

### Pol I inhibition perturbs splicing programs

These findings raised the possibility that Pol I activity relates to splicing programs. To assess global transcriptional response to Pol I inhibition stress, we conducted RNA-seq analysis in p53^+/+^ and p53^-/-^ cells and A375 melanoma cells (wild-type for both *TP53* and *RPL22*) following treatment with BMH-21 (**Figure S5A**). The alterations of Pol II transcripts were broad and robust, and as expected, GSEA pathway analyses showed prominent activation of p53 pathway in the p53 proficient cells, and increased expression of p53 target genes (**Figure S5B**). We then conducted rMATS analyses of the RNA-seq data and found 300 - 600 splicing events in each line using stringent criteria (FDR <0.05, minimum 10 reads for each event) (**Figures 6A–6C**, **Figure S5C** and **Tables S7–S9**). Spliceosome was the most significant pathway identified (**Figures 6A–6C**). This finding was striking, as the splicing factors undergo autoregulation (Lareau et al. 2007). By analyzing shared splicing factor alterations in the p53^+/+^ and p53^-/-^ cells we saw many splicing factors that were repressed following Pol I stress (**Figure 6D**). These unbiased analyses demonstrated that Pol I inhibition stress perturbs splicing and splicing factor expression.

**Figure 6.**
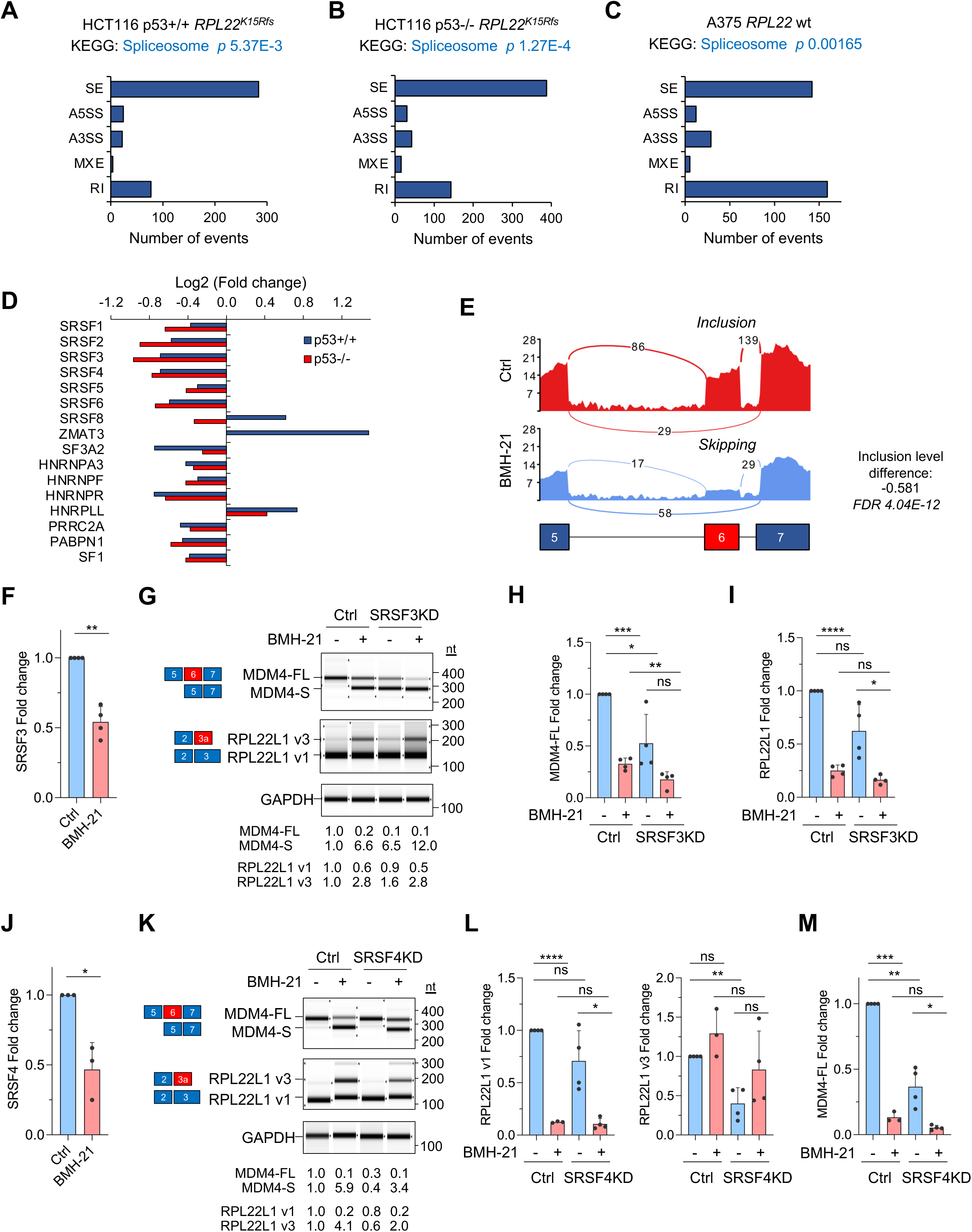
Pol I inhibition perturbs splicing programs. (**A-C**) rMATS and pathway analyses in BMH-21 treated cells. **A**, p53^+/+^ cells, **B**, p53^-/-^ cells and **C**, A375 melanoma cells. (**D**) RNA-seq data of splicing factors. n = 3 biological replicates. Statistics, q-values < 0.05. (**E**) Sashimi plot of MDM4 splicing following BMH-21 treatment in A375 cells. (**F**) qPCR for SRSF3 following BMH-21 treatment. (**G**) MDM4 and RPL22L1 isoforms following SRSF3 knockdown. Representative sqPCR of n = 3 is shown. (**H**) qPCR for MDM4 isoforms in parent and SRSF3^KD^ cells. (**I**) qPCR for RPL22L1 in parent and SRSF3^KD^ cells. (**J**) qPCR for SRSF4 following BMH-21 treatment. (**K**) MDM4 and RPL22L1 isoforms in SRSF4^KD^ cells. Representative sqPCR of n = 2 is shown. (**L**) qPCR for RPL22L1 variants in parent and SRSF4^KD^ cells. (**M**) qPCR for MDM4-FL in parent and SRSF4^KD^ cells. Data are representative of at least three independent experiments (F, G, H, I, J, L, M) and mean±SD are shown. *p* values, Student’s two-sided t-test. *ns*, non-significant, * <0.05, ** <0.01, *** <0.001, **** <0.0001. See also Figure S5, Tables S7, S8 and S9.

We then focused on identification of splicing factors involved in the splicing of MDM4 and RPL22L1. Isoform analysis showed that BMH-21 decreased the expression of MDM4-FL in all cell lines (**Figure S5D**). We visualized MDM4 splicing using Sashimi plot, which showed that BMH-21- treatment significantly increased exon 6 skipping (**Figure 6E**). The previously identified MDM4 splicing factors are serine-arginine rich splicing factor 3 (SRSF3), protein arginine methyltransferase 5 (PRMT5), and p53 target gene ZMAT3 (Bezzi et al. 2013; Dewaele et al. 2016; Gerhart et al. 2018; Bieging-Rolett et al. 2020). SRSF3 and PRMT5 promote MDM4 exon 6 inclusion and ZMAT3 exon 6 skipping. ZMAT3 increases MDM4-S expression in a p53-dependent manner (Bieging-Rolett et al. 2020). ZMAT3 was induced by BMH-21 but only in p53-proficient cells (**Figure 6D**). Therefore, ZMAT3 is unlikely to be responsible for the p53-independent regulation of MDM4. Instead, SRSF3 was repressed in all cell lines (**Figure 6D**). We further analyzed the effect of BMH-21 on SRSF3 by using qPCR and found that SRSF3 transcript was downregulated by BMH-21 by 50%, and its protein abundance by 40% (**Figure 6F** and **Figure S5E**). We then used sqPCR and qPCR to detect MDM4 splicing isoforms and saw that SRSF3 knockdown decreased basal MDM4-FL, which was further exacerbated by BMH-21 (**Figures 6G** and **6H** and **Figure S5F**). These approaches confirm that SRSF3 regulates MDM4 exon 6 inclusion in a Pol I activity-dependent manner. However, SRSF3 did not alter RPL22L1 splicing or affect that by BMH-21 (**Figures 6G** and **6I**).

SRSF4 has been implicated earlier to alter RPL22L1 splicing (Larionova et al. 2022). As expected based on RNA-seq, BMH-21 decreased SRSF4 (**Figure 6J**). We then used SRSF4 KD and observed that it significantly decreased RPL22L1 variant 3 expression (**Figures 6K** and **6L** and **Figure S5G**). Interestingly, the knockdown also decreased MDM4-FL (**Figure 6M**). We conclude that multiple SRSF splicing factors are regulated by Pol I inhibition and alter the splicing of MDM4 and RPL22L1.

### RPL22 connects rRNA synthesis activity with splicing

RPL22 is a constituent member of the ribosome and binds 28S rRNA. This interaction is well resolved in several ribosome cryo-EM structures and mediated by RPL22 codons identified critical for RNA binding (Houmani and Ruf 2009; Natchiar et al. 2017; Faille et al. 2023; Holm et al. 2023). RPL22 also interacts with cellular and viral RNAs that have a small hairpin stem loop with a loosely defined sequence (Toczyski et al. 1994; Dobbelstein and Shenk 1995). In zebrafish, RPL22 negatively regulates RPL22L1 by binding to a stem loop on exon 2 (O’Leary et al. 2013). We identified two potential stem loops in human RPL22L1 intron 2 (**Figure S6A**). Strikingly, these sequences overlapped the RPL22L1 variant 3 intronic splice site. Given this we hypothesized that the ability of RPL22 to bind to RPL22L1 pre-mRNA mediates the repression of RPL22L1. To test whether the intronic sequences are essential for RPL22L1 repression, we expressed RPL22L1 from a cDNA vector. As shown in **Figure S6B**, BMH-21 prominently decreased the endogenous but not the ectopically expressed RPL22L1 transcript.

Comparative genome-wide RNA-binding by RPL22 and RPL22L1 has not been analyzed previously. To assess this, we generated cell lines that stably express RPL22 and RPL22L1 HaloTag fusion proteins, whereas cells expressing HaloTag served as control (**Figure S6C**). We used GoldCLIP-seq (Gu et al. 2018) for RNA-pulldowns that enable highly effective RNA-interactome analyses by the virtue of covalent coupling to HaloTag resins and the precision on binding site analysis gained by using RNAse digestion (**Figure 7A**). RPL22 and RPL22L1 interactomes included several RNA species, most prominently rRNA and mRNAs but also a large number non-coding RNAs such as long intergenic non-coding RNAs, miRNAs, small nucleolar RNAs, and RNAs involved in major and minor site splicing (**Figure S6D** and **Tables S10** and **S11**). RPL22-HaloTag showed strong preferential binding to 28S rRNA sequence in region III identified in ribosome structural studies as the RPL22 interface (**Figure 7B** and **Fig. S6E**) (Natchiar et al. 2017; Faille et al. 2023; Holm et al. 2023). RPL22L1-HaloTag was also bound to this site, albeit at greatly lower frequency, likely reflecting RPL22 and RPL22L1 stoichiometry as ribosome constituents. This 28S rRNA sequence has a long stem structure with two mirrored loops but lacks the small stem loops (**Figure 7B**). We did not detect binding of either protein to 28S rRNA sequences previously identified by SELEX analysis (Dobbelstein and Shenk 1995).

**Figure 7.**
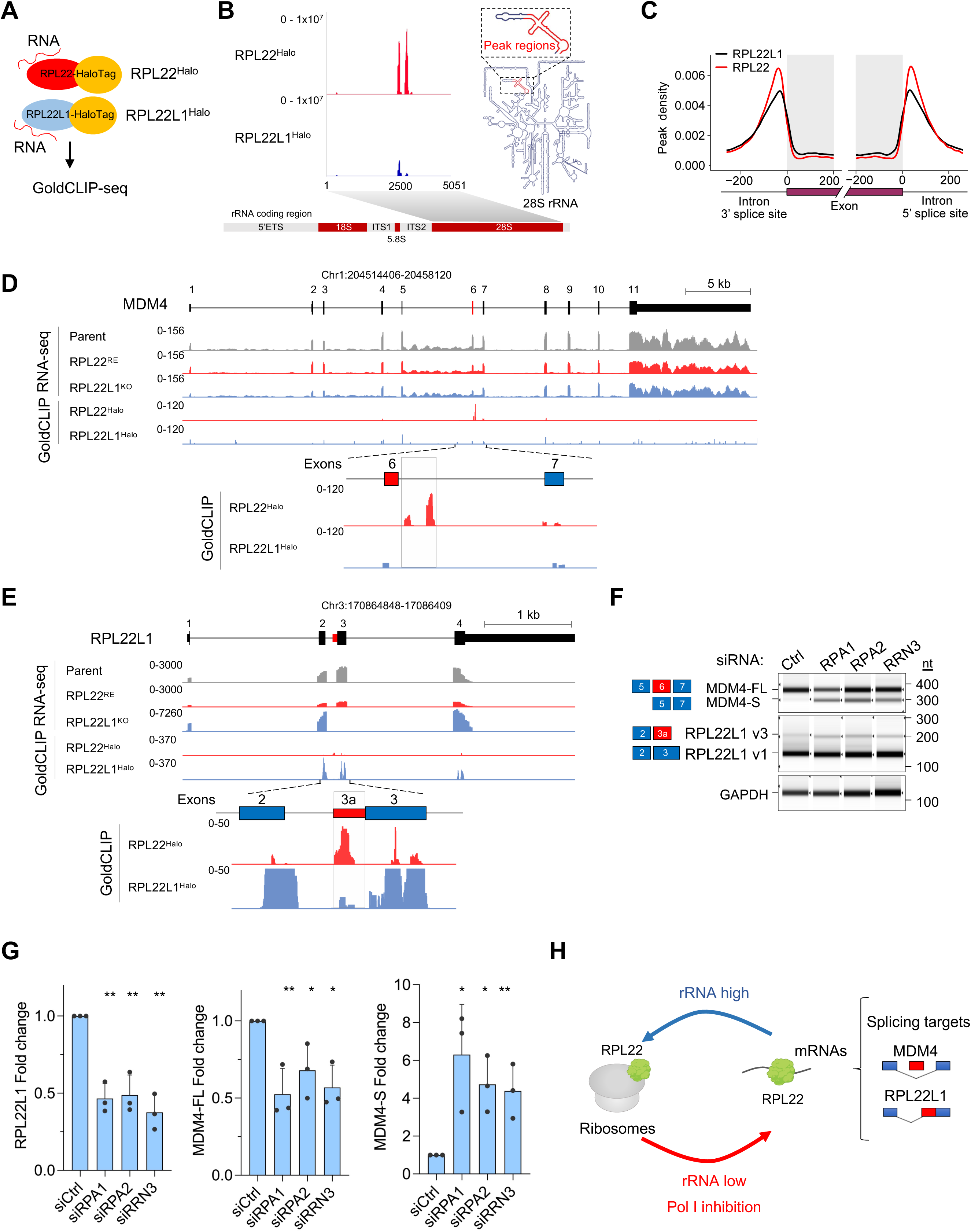
RPL22 coordinates splicing with rRNA synthesis. (**A**) Schematic of GoldCLIP-seq. (**B**) RPL22^Halo^ and RPL22L1^Halo^ binding to 28S rRNA. (**C**) Metagene plots of RNA binding peaks in RPL22^Halo^ and RPL22L1^Halo^ cells. (**D**) MDM4 expression tracks and HaloTag-fusion protein binding peaks of RPL22 and RPL22L1. Bottom tracks visualize binding peaks in intron 6. (**E**) RPL22L1 expression tracks and HaloTag-fusion protein binding peaks of RPL22 and RPL22L1. Bottom tracks visualize binding peaks in intron 2/exon 3a. (**F**) RPL22L1 and MDM4 isoforms of Pol I knockdown cells. Representative sqPCR of n = 3 is shown. (**G**) qPCR for RPL22L1 and MDM4 isoforms in Pol I knockdown cells. n = 3 independent biological replicates, mean±SD. (**H**) Schematic model of connection of rRNA synthesis to splicing by RPL22. *p* values, Student’s one-sided t-test. *ns*, non-significant, * <0.05, ** <0.01, *** <0.001, **** <0.0001. See also Figure S6, Tables S10 and S11.

The distribution of RPL22 and RPL22L1 on coding genes was striking as both were enriched on introns (**Figure S6F**). Metagene analysis on read coverage showed their enrichment at 5’ and 3’ splice site junctions (**Figure 7C**). Motif analyses using HOMER identified several significant motifs most of which were distinct for RPL22 and RPL22L1 (**Figure S6G**). Strikingly, RPL22-HaloTag was prominently and exclusively enriched on MDM4 intron 6 as two distinct peaks that overlapped the predicted SRSF3 and ZMAT3-binding sites (**Figure 7D**) (Dewaele et al. 2016; Bieging-Rolett et al. 2020). Instead, RPL22L1 had low read coverage on MDM4 pre-mRNA (**Figure 7D**). RPL22, but not RPL22L1-HaloTag was enriched on RPL22L1 intron 2 overlapping the splice site for variant 3 (**Figure 7E**). Interestingly, RPL22L1-HaloTag was enriched on its own exons, suggesting that it could regulate the stability of its transcript (**Figure 7E**).

We conducted an intersection analysis to ask whether RPL22 or RPL22L1 binding peaks overlapped with the splicing alterations detected in RPL22^RE^ and RPL22L1^KO^ cells. We identified approximately 1,000 genes each for RPL22 and RPL22L1 where splicing events overlapped their binding sites (**Figure S6H**). Of these, nearly 300 genes were shared, such as MDM4, CD44, MLH1, MLH3 (**Figure S6H**). Despite substantial overlap of their binding sites, hundreds of genes were unique suggesting independent regulatory activities by the paralogs (**Figure S6H**). For RPL22-bound transcripts, the majority were represented by exon skipping and mixed exon splicing events (**Figure S6I**).

To further assess the dependency of the splicing events on rRNA synthesis activity we used genetic inhibition of Pol I. We knocked down Pol I enzyme catalytic subunits RPA1, RPA2 and the essential preinitiation factor RRN3. Each knockdown decreased Pol I transcription by half (**Figures S6J** and **S6K**). The splicing of both MDM4 and RPL22L1 substantially and significantly altered (**Figures 7F** and **7G**). MDM4-FL was decreased by 40% and MDM4-S was increased by 5-fold by Pol I knockdown (**Figure 7G**). Furthermore, RPL22L1 variant 1 decreased whereas RPL22 was unaffected (**Figure 7G** and **Figure S6L**). Consistent with the drug effect, genetic inhibition of Pol I also significantly decreased SRSF3 expression (**Figure S6M**). We conclude that both chemical and genetic inhibition of Pol robustly alter the expression of MDM4 and RPL22L1 isoforms.

These data are consistent with a model that RPL22 negatively regulates the splicing and expression of RPL22L1 and MDM4 and that this depends on rRNA synthesis activity (**Figure 7H**). Consequently, RPL22 coordinates rRNA synthesis and splicing.

## Discussion

Here we describe a ribotoxic stress mechanism that connects rRNA synthesis to cellular splicing by ribosomal protein RPL22. We find that cancer-haploinsufficient RPL22, a constitutive member of the 60S ribosome and 28S rRNA interacting ribosomal protein, binds to hundreds of mRNAs affecting their splicing and is essential for the expression and splicing of MDM4 and its paralog RPL22L1. Strikingly, the ability of RPL22 to repress the expression of RPL22L1 and MDM4 is dependent on rRNA synthesis activity. We present a model (**Figure 7H**) of how the rRNA synthesis activity defines the distribution of RPL22 between the ribosomes and the target pre-mRNAs controlling the expression of transcripts such as RPL22L1 and the p53 repressive isoform MDM4-FL. Whereas high rRNA synthesis activity sequesters RPL22 into the ribosomes, therapeutic inhibition of Pol I restores RPL22 interaction with its mRNA targets. Given that RPL22 binds mRNAs at exon-intron junctions and changes the splicing of hundreds of genes, it suggests that the impact of these regulatory events in *RPL22*-altered cancers is broad. These findings reveal an unexpected coordination of cellular splicing with rRNA synthesis mediated by RPL22.

Cancer cell line vulnerability screens identified that the sensitivity to Pol I inhibition is linked with *RPL22* mutation and high expression of RPL22L1 and MDM4. MSI cancer cell lines were especially sensitive to the inhibitors. Cell-based competition models validated that RPL22, RPL22L1 and MDM4 each conveyed sensitivity to the Pol I inhibitors. Earlier studies had recognized RPL22 and RPL22L1 synthetic paralog relationship and high frequency of *RPL22* mutations in MSI cancers that also highly express RPL22L1 and MDM4-FL (O’Leary et al. 2013; McDonald et al. 2017; Ghandi et al. 2019). These markers were proposed to mediate the sensitivity to MDM2 inhibitor Nutlin-3a, but this notion was not further explored (Ghandi et al. 2019; Weinstein et al. 2023). We dissected the identified sensitivities and their mechanistic underpinnings and found that pharmacological and genetic inhibition of Pol I decreased the expression of RPL22L1 and MDM4, and that this decrease coupled with their splicing alterations in a RPL22-dependent manner. Nutlin-3a lacked these activities. This mechanism is distinct from, but parallel to the transcription-stress induced reorganization of the nucleolus and inhibition of MDM2 by ribosomal proteins RPL5 and RPL11. Both mechanisms uniquely converge on p53. MDM2 inactivation stabilizes p53 while MDM4 splicing change abrogates p53 transcriptional repression. In response to Pol I inhibition, both pathways are activated and convey robust activation of p53. Intriguingly, the RPL5/MDM2 complex also incorporates SRSF1, which is strengthened by Pol I inhibition to activate p53 (Gu et al. 2018). However, MDM4 also has p53-independent oncogenic activities, and RPL22L1 is a putative oncogene, suggesting that their repression is therapeutically valuable independent of p53 (Gembarska et al. 2012; Rao et al. 2012; Klein et al. 2021; Ueda et al. 2021). As we have demonstrated earlier, and show here, *TP53* is nonessential for Pol I inhibition (Peltonen et al. 2014b).

Earlier studies found that Pol I stress regulated the stability of MDM4 (Gilkes et al. 2006; Li and Gu 2011). However, here we uncover a mechanism how Pol I inhibition switches MDM4 splicing by increasing exon 6 skipping leading to the expression of an NMD-targeted MDM4-S isoform and loss of protein expression. The splicing change may involve several splicing factors including SRSF3 and ZMAT3 previously identified to increase exon 6 inclusion and skipping, respectively (Dewaele et al. 2016; Bieging-Rolett et al. 2020). RPL22 is a tumor suppressor, but it is not known how it conveys this function (Rao et al. 2012; Rao et al. 2016). *TP53* mutations are strikingly underrepresented in *RPL22* mutant tumors. The highly frequent *RPL22* mutations and haploinsufficiency in MSI are suggestive that these alterations predispose to not only increased expression of MDM4-FL and repression of p53, but elevated RPL22L1. A decrease in RPL22L1 expression decreases colon cancer cell growth as shown here and by Rao et al. (Rao et al. 2019) and leads to large-scale changes in gene expression and splicing. The expression of RPL22L1 variant 3 was previously detected in glioblastoma neurospheres and its splicing was promoted by low pH, hypoxia and SR-splicing factors (Larionova et al. 2022). As variant 3 lacks the RNA interaction domain of RPL22L1, it is unclear whether it retains RNA-binding function. Notably, acidosis and hypoxia compromise Pol I transcription (Mekhail et al. 2006) suggesting that the mechanism identified here could lead to altered RPL22L1 splicing. Our analysis is limited to the detection of RNA transcripts as we are unable to confirm the expression of RPL22L1 variant 1 or variant 3 proteins due to lack of antibodies with sufficient specificity and sensitivity.

Effects of RPL22 and RPL22L1 on splicing in cancers have not been previously detailed. By residing at the splicing junctions, they may fine-tune the coordinated activity of several splicing factors. The comprehensive RPL22 and RPL22L1 expression, splicing, interactome and motif analyses identified, despite some notable shared targets such as MDM4, mostly distinct regulatory targets. The RNA-interactome analyses showed binding of RPL22 to introns of both MDM4 and RPL22L1, and while the RPL22L1 intron 2 had two RPL22-interacting stem loops precisely at the splice site, the MDM4 intron did not. In contrast, RPL22L1 lacked discernible binding to these sites, suggesting that their regulatory roles in cancer cells are distinct. Instead, RPL22L1 is prominently bound to its own exons, implicating autoregulation of the stability of its mRNA. Future studies should explore the scope of their impact on both splicing and regulation of RNA stability.

These markers for Pol I inhibitor sensitivity are frequently altered in cancers and facilitate future development of tumor-agnostic therapeutic strategies. In addition to *RPL22* mutations in dMMR cancers of the colon, stomach and endometrium, *RPL22* is mutated in 10% T-cell acute lymphoma/leukemia and *RPL22* locus at chromosome 1p36 is frequently deleted in sporadic colorectal cancers leading to the overexpression of RPL22L1 and 5-FU resistance (Rao et al. 2012; Rao et al. 2019). *RPL22L1* 3p26 locus is amplified in ∼30% squamous lung cancers and over 20% ovarian and esophageal cancers (cBio Portal) and MDM4 is highly expressed in melanoma, retinoblastoma and acute myeloid leukemia (Laurie et al. 2006; Gembarska et al. 2012; Marine and Jochemsen 2016; Ueda et al. 2021). Consequently, we show that Pol I inhibition is highly effective in xenograft, syngeneic and PDX models with alterations in *RPL22* and MDM4. WRN, and more recently ATR, has been identified as a synthetic lethality gene in MSI, and WRN inhibitors such as HRO761 have entered phase 1 clinical trials (Chan et al. 2019; N et al. 2020; Wang et al. 2023; Zong et al. 2023; Ferretti et al. 2024). However, these inhibitors are effective only in MSI cancers with high repeat expansion signatures (Morales-Juarez and Jackson 2022). They are thus mechanistically distinct from the Pol I inhibitors, which have a broader vulnerability profile. Immune checkpoint inhibitors have become standard-of-care in many dMMR tumors, and their effectiveness is encouraging although ICI resistance remains a barrier (Jin and Sinicrope 2022). Here, we find that the Pol I inhibitors improved the efficacy of ICI and increased T-cell infiltration in the tumors while reducing the presence of myeloid cells. The combinatory effectiveness of the Pol I inhibitors with ICI is promising but requires further testing. A high mutational load is considered to drive the recognition of the tumor by immune cells (Mandal et al. 2019). Alternative splicing is a source of tumor antigenicity and generates neoepitopes more frequently than by mutation burden and thus contributes to the immune response (Frankiw et al. 2019; Dersh et al. 2021; Lu et al. 2021). Given that *RPL22* mutation and haploinsufficiency is highly prevalent in MSI it is reasonable to propose that splicing alterations contribute to altered tumor antigenicity and impart ICI responsiveness. Future studies should test this possibility.

We propose that RPL22 coordinates rRNA transcription and splicing of mRNA genes and that the cancer splicing program is rewired due to *RPL22* genetic alterations. We predict this circuitry is critical during times of transcriptional bursts, such as growth, developmental processes and pathological states such as cancer. In other words, Pol I transcriptional state is coordinated with systemic post-transcriptional regulation of splicing programs.

## Acknowledgment

We thank all Laiho lab past members for support and discussions. We thank the Maryland Advanced Research Computing Center and the following Johns Hopkins University core facilities for expert technical assistance and services: Deep Sequencing Core, Experimental and Computational Genomics Core, Genetics Resources Core Facility, Analytical Pharmacology Shared Resource, Flow Cytometry Development Core, Oncology Tissue Services, and Animal Resources. This work was funded by grants from NIH (R01 GM121404, P30 CA006973), Blue One Biosciences LLC, Commonwealth Foundation, Mary Kay Ash Charitable Foundation, Academy of Finland (288364), Maryland Cigarette Restitution Fund, and Harrington Scholar-Innovator Award (to M.L.). W.F. is supported by NCI K99 CA279786.

## Author contributions

Conceptualization, M.L.; methodology, W.F., H.L., T.E.D., G.E., W.C.R., B.O., M.A.R., N.V.R.; investigation, W.F., H.L., G.C.S., A.B., C.E.D., T.D., W.C.R., Q.Z., B.O., N.V.R.; writing - original draft, M.L.; writing and editing, W.F., H.L., N.V.R.; writing – review and editing, all authors; supervision, A.M.D., J.C.B., M.L.; funding acquisition, W.F., M.L.

## Materials and methods

### Materials availability

Plasmids, cell lines, and reagents generated in this study are available upon request to the corresponding author.

### Data and code availability

RNA, ChIP and Gold-CLIP sequencing data have been deposited to GEO with the following identifiers GSE267499, GSE267500, GSE267501, GSE267502, GSE267503.

### Cell lines

Cell lines used in this study are detailed in **Methods Table 1** and were maintained as indicated at 37°C and 5% CO_2_ in a humidified atmosphere. Authentication of cell lines was conducted using short tandem repeat profiling at the Johns Hopkins University Genetic Resources Core Facility using GenePrint 10 (Promega, cat # B9510).

**Methods Table 1.** Cancer cell lines, genotypes and sources.

| Cell line | Source# | Lineage | Gender | MSI | TP53 | RPL22<sup>K15Rfs</sup> | STR verified | Cell culture medium* |
| --- | --- | --- | --- | --- | --- | --- | --- | --- |
| 22RV1 | ATCC | Prostate | male | unstable | mut | mut | yes | RPMI-1640+10% FBS |
| A375 | ATCC | Melanoma | female | stable | wt | wt | yes | DMEM+10% FBS |
| DLD1 | F. Bunz | Colorectal | male | unstable | mut | mut | yes | DMEM+6% FBS |
| Gp2D | Sigma | Colorectal | female | unstable | wt | mut |  | DMEM+10% FBS |
| HCT-8 | ATCC | Colorectal | male | unstable | wt | wt | yes | RPMI-1640+5% FBS |
| HCT116 | F. Bunz | Colorectal | male | unstable | wt | mut | yes | McCoy's 5A+10% FBS |
| HEC151 | JCRB | Endometrial | female | unstable | mut | mut | yes | EMEM+15% FCS |
| HEC265 | JCRB | Endometrial | female | unstable | wt | mut |  | EMEM+15% FCS |
| IM95 | JCRB | Gastric | male | unstable | wt | mut | yes | DMEM+10% FBS |
| LNCaP | ATCC | Prostate | male | stable | wt | mut | yes | RPMI-1640+10% FBS |
| LoVo | ATCC | Colorectal | male | unstable | wt | mut |  | F-12K+10% FBS |
| SNU-1 | ATCC | Gastric | male | unstable | wt | mut |  | RPMI-1640+10% FBS |
| NUGC-3 | JCRB | Gastric | male | unstable | mut | wt |  | RPMI-1640+10% FBS |

### Chemical compounds

12H-Benzo[g]pyrido[2,1-b]quinazoline-4-carboxamide, N-[2(dimethylamino)ethyl]-12-oxo (BMH-21) and BOB-42 were synthesized as described in Colis et al. (Colis et al. 2014a) and verified for purity using liquid chromatography/mass spectrometry (LC/MS) and ^1^H nuclear magnetic resonance (NMR) spectroscopy. The following compounds were purchased from commercial sources: Actinomycin D (Sigma-Aldrich, cat # A1410), Nutlin-3a (Sigma-Aldrich, cat # SML0580), PRMT5 inhibitor GSK3326595 (MedChemExpress, cat # HY-101563), CLK1 inhibitor TG003 (MedChemExpress, cat # HY-15338). Where indicated, cells were treated with actinomycin D (5 nM), Nutlin-3a (10 µM), GSK3326595 (0.5 µM) and TG003 (100 µM).

### Vectors and vector constructions

pCMV6-RPL22L1-myc-DDK vector was obtained from Origene (cat # RC211790). Human RPL22 and RPL22L1 cDNAs were amplified from pLenti-GIII-CMV-RPL22-HA and pLenti-GIII-CMV-RPL22L1-HA (Applied Biological Materials, cat # LV291610, cat # LV802168) then cloned into a PB-EF1α-MCS-IRES-GFP PiggyBac expression vector (System Biosciences, cat # PB530A-2). HaloTag (Los et al. 2008) was amplified from pSMART-Rps3-HaloTag7-3xFlag (gift from Heeseon An) and P2A-puro was amplified from lentiCRISPRv2 puro (Addgene, cat # 98290). The amplified PCR products were cloned using GeneArt™ Seamless Cloning and Assembly Enzyme Mix (Invitrogen, cat # A14606) according to the manufacturer’s protocol to generate PB-EF1α-RPL22-HaloTag-P2A-puro and PB-EF1α- RPL22L1-HaloTag-P2A-puro. Super PiggyBac Transposase Expression Vector (System Biosciences, cat # PB210PA-1) was transfected with PB-EF1α-RPL22-HaloTag-P2A-puro and PB-EF1α-RPL22L1- HaloTag-P2A-puro to efficiently integrate the inserts into the genome. For GFP competition assay, pLKO.1-puro-CMV-TurboGFP plasmid (Sigma-Aldrich, cat # SHC003) was used to generate stable cell lines expressing GFP. Vectors were confirmed by Sanger sequencing.

### Cancer cell line analyses

Cancer cell line screening on the Oncosignature panel of lines detailed in **Supplementary Tables 1 and 2** was performed at Horizon Discovery Limited (Cambridge, UK). Cells were thawed from liquid nitrogen, expanded and once dividing at their expected doubling times plated in their respective growth media in black 384-well tissue culture plates at 500 to 1500 cells/well and equilibrated via centrifugation. 24 hours after the plating, a “time zero” set of assay plates (V0) was collected and ATP levels were measured using CellTiterGlo2.0 (Promega) using Envision plate readers (PerkinElmer).

The screen was conducted for each drug using 10-point 3-fold dilution series starting at 10 µM in triplicate. Cell viability was measured using CellTiterGlo after 72 hours of incubation, and the response curves were used to determine GI_50_ using a Horizon Discovery proprietary software. GI was defined as in (Shoemaker 2006) as 100 x [1- (T - V0)/V-V0], where V0 is the vehicle-treated level at time zero, T the treated level and V the vehicle level at 72 hrs. Z-factors for each plate were determined, and assay plates were accepted if the following quality control standards were met: consistent relative luciferase values throughout the entire experiment, Z-factor scores >0.6, and consistent behavior of untreated/vehicle controls on the plate.

### Cell line correlative analyses

RNA expression, protein array (RPPA), mutations (WES), copy number variation, metabolomics, histone modifications, promoter methylation and microRNA omics data were obtained from DepMap (19Q3) (Tsherniak et al. 2017). CRISPR gene dependency scores (22Q2) were downloaded from DepMap portal. Batch correlation between BMH-21 and BOB-42 GI_50_ and -omics data were performed in the R. Spearman method, and statistics were used in the batch correlations. The *p* values were computed and then converted to the negative logarithm of the *p* value for plotting.

### qPCR and semiquantitative PCR

RNA was extracted from cells using RNeasy Mini Kit (Qiagen). RNA was reverse transcribed using SuperScript II Reverse Transcriptase (Invitrogen). To perform qPCR, the resulting cDNA was mixed with PowerUp SYBR Green Master Mix (Applied Biosystems by Thermo Fisher Scientific) and the appropriate primer pairs. Analyses were conducted in triplicate using BioRad CFX384 Real-Time System – C1000 Touch Thermal Cycler or the Applied Biosystems QuantStudio 12K Flex Real-Time PCR System. All results were normalized to GAPDH, and RNA levels were quantified using the ΔΔCt method. All PCR reactions were conducted in triplicates, and the data presented represent a minimum of three independent biological replicates. Bar graphs represent the relative ratios of each gene to GAPDH values and control sample.

For semiquantitative PCR, 20 ng cDNA was used in a 10 µL PCR reaction with Q5 Hot start 2x high-fidelity PCR master mix (New England Biolabs, cat # M0494s). After 21 cycles for GAPDH and 26 cycles for other transcripts, 1 µL was mixed with 3 µL D1000 sample buffer (Agilent Technologies, cat # 5190-6502) and run on Agilent 4200 TapeStation platform with D1000 screenTape (Agilent Technologies, cat #5067-5582).

The following primer pairs were used:

MDM4-FL (forward 5’GATGCTGCTCAGACTCTCGC, reverse 5’TGCACTTTGCTTCAGTTGGTC) MDM4-S (forward 5’GCCACTGCTACTACAGCAAAG, reverse 5’TCTGAGGTAGGCAGTGTGGG) MDM4 semi-qPCR (forward 5’AGGTGCGCAAGGTGAAATGT, reverse 5’TCTGAGGTAGGCAGTGTGGG) CDKN1A (forward 5’TTAGCAGCGGAACAAGGAGT, reverse 5’TCAACGTTAGTGCCAGGAAA) MDM2 (forward 5’CGGAAAGATGGAGCAAGAAG, reverse 5’GCGCTCGTACGCACTAATC) RPL22 (forward 5’TGACATCCGAGGTGCCTTTC, reverse 5’GTTAGCAACTACGCGCAACC) RPL22L1 (forward 5’GATTGGCTTCGAGTGGTTGC, reverse 5’CCCTGTAAGGGGAGCCTTTG) RPL22L1myc (5’CTTCGAGTGGTTGCATCTGA, reverse 5’CAGATCCTCTTCTGAGATGAGTTTC) RPL22L1 variant 1 (5’AGACAGGAAGCCCAAGAGGT, reverse 5’TCCCGAGATTTCCAGTTTTG) RPL22L1 variant 3 (5’GGGAAGAGAACCCTAGATATTCA, reverse 5’TCCCGAGATTTCCAGTTTTG) GAPDH (forward 5’GGCCTCCAAGGAGTAAG ACC, reverse 5’AGGGGTCTACATGGCAACTG) RPA1 (forward 5’GCGTGGTGACTCCGGGCTTG, reverse 5’CAGGCCGTTTGCCGATGGGT) RPA2 (forward 5’GCCTCGCGGTGCAGGCTATAC, reverse 5’CCTTCGGCCCCGGCATTCTG) RRN3 (forward 5’CGGTTTGGTGGAACTGTGAC, reverse 5’AGAACGGAATTCTAGCAGCCA) SRSF3 (forward 5’AATAAAGGCGAGGAGAAGGCG, reverse 5’CACACGCTCCGATGAGTCTT) SRSF4 (forward 5’CAGCCATCACTGCCGTTG, reverse 5’CGCAGATCATCAAACTCCACA) 28S (forward 5’TGGGTTTTAAGCAGGAGGTG, reverse 5’AACCTGTCTCACGACGGTCT) 5’ETS (forward 5’GAACGGTGGTGTGTCGTT, reverse 5’GCGTCTCGTCTCGTCTCACT).

### Immunoblotting

Cells were lysed in RIPA lysis buffer (Millipore Sigma, cat # 20-188) supplemented with protease inhibitors (Millipore Sigma, cat # 4693132001). Protein concentrations were determined using Pierce BCA Protein Assay Kit (Thermo Fisher Scientific, cat # PI23225). An equal amount of protein per sample was mixed with Laemmli sample buffer (Bio-Rad), DTT, boiled and run on an NuPAGE 4-12% Bis-Tris gel (Invitrogen, cat # NP0322BOX) and transferred to Immobilon PVDF (0.45 μm) membrane (MilliporeSigma, cat # IPVH00010). The membrane was blocked with 5% milk followed by primary and secondary antibodies. The primary antibodies used were RPL22 (Abcam, cat # ab229458), MDM4 (Bethyl Laboratories, cat # A300-287A), SRSF3 (Thermo Fisher Scientific, cat # 33-4200), p53 (DO1, Thermo Fisher Scientific, cat # MA512571), anti-HaloTag (Promega, cat # G9211), α-Actin (Abcam, cat # ab5694), α-Tubulin (Abcam, cat # ab7291), and GAPDH (Abcam, cat # ab8245). Horseradish peroxidase (HRP)-conjugated secondary antibodies were from Dako (cat # P026002-2 and cat # P021702-2) and detected using either SuperSignal West Pico PLUS Chemiluminescent Substrate (Thermo Fisher Scientific, cat # SKU:34579) or Western Lightning Plus-ECL Enhanced Chemiluminescence Substrate (Perkin Elmer, cat # NEL103001EA). The membranes were imaged using the BioRad Molecular Imager ChemiDoc XRS+ System. Protein densitometry was conducted using Image Lab Software.

### Cell counting and competition assays

Isogenic HCT116 cells were transfected with RPA1 siRNAs, incubated for 48 hours and treated with or without BMH-21 for 48 hours. Cells were then counted using Cellometer AutoT4. For competition assay, stable GFP-expressing HCT116 p53^+/+^ cells were generated, and 0.75 million each of the GFP- parent and stable knockdown cells were mixed and cultured for 2 weeks in the presence of absence of BMH-21 (0.25 µM). Cells were then trypsinized and analyzed by flow cytometry using Beckman Coulter CytoFLEX S 3-laser flow cytometer.

### Gene silencing and knockouts

Cells were transfected using Lipofectamine RNAiMax (Invitrogen, cat # 13778075) transfection reagent and incubated for 48 hours. siRNAs were obtained from Ambion, Thermo Fisher Scientific as follows. Negative control siRNA (cat # AM4635), RPA1 cat # s403; RPA2 cat # s38603; RRN3 cat # s29324; SRSF3, cat # 12731. Stable shRNA gene knockdowns were generated as in Pitts et al.^21^ Vectors for shRNAs were obtained from Sigma-Aldrich as follows. RPL22, TRCN0000075015; RPL22L1, TRCN0000247704; MDM4, TRCN0000003859; SRSF4 TRCN0000231448. RPL22L1 knockout was carried out using Gene Knockout Kit v2 (Synthego). To achieve high knockout efficiencies, two guide sgRNAs targeting RPL22L1 exon 3 were designed and synthesized by Synthego: sgRNA1: 5’GGAGCAATTTCTACGGGAGA, sgRNA2: 5’GAAACAGTTCTCTAAAAGGT.

Cas9 and sgRNA ribonucleoprotein (RNP) complexes were transfected to HCT116 p53^-/-^ cells using nucleofection according to the Synthego protocol. Briefly, sgRNA (30 µM) and 1 µL Cas9 were mixed with 18 µL Nucleofector™ Solution and incubated for 10 minutes at room temperature. Cells were then suspended in buffer SE and transfected using 4D-Nucleofector (Lonza) program EN-113. 72 hours post nucleofection, single cells were seeded in 96-well plates using limiting dilution. After clonal expansion, single clones were picked followed by genotyping and Sanger sequencing. For Sanger sequencing the following primers were used: forward 5’GGGTTCTCTTCCCTAGTTTTGC (intron 2) and reverse 5’TTTCCCATTTATGTAGGCCTTT (intron 3). The knockout of exon 3 was further confirmed by qPCR.

### RNA-seq

As indicated, cells were treated with DMSO or BMH-21 (1 µM) for 6 hours. Three biological replicates were prepared for each treatment. RNA was extracted with TRIzol (Thermo Fisher Scientific, cat # 15596026). RNA-seq library for Illumina platform sequencing was prepared using Illumina TruSeq stranded total RNA Sample kit following manufacturer’s protocol followed by sequencing on Illumina NextSeq 500 for 2X75 bp paired-end reads. Reads were analyzed using Tophat 2 and Cuffdiff 2.0, respectively for alignment to reference genome and differential expression detection. Pathway analyses were conducted using Gene Set Enrichment Analysis (GSEA). RNA-seq of HCT116 parent, RPL22^RE^ and RPL22L1^KO^ cells was conducted on Illumina NovaSeq 6000. Paired-end reads were mapped using HISAT2 software and StringTie was used to estimate transcript abundances and differential expression. Significantly differentially expressed transcripts were defined using a p-value cutoff < 0.05.

### Alternative splicing analysis

Alternative splicing analysis was conducted using rMATS 4.1.1 (Shen et al. 2014) based on triplicate RNA-seq samples. Splicing differences were calculated for five classes of alternative splicing events, skipped exons (SE), alternative 5’ splice sites (A5SS), alternative 3’ splice sites (A3SS), mutually exclusive exons (MXE), and retained introns (RI). Significant differential alternative splicing events were defined as FDR (false discovery rate) < 0.05 with a requirement of an average read count > 10 per event. Data is reported on reads covering exon junctions. Sashimi plot visualization of alternative splicing events was generated by the rmats2sashimiplot function in rMATS. Inclusion level differences are reported and represent the average difference between the splicing events calculated from normalized counts.

### GoldCLIP-seq

GoldCLIP-seq was performed according to protocol designed by Gu et al. (Gu et al. 2018) with modifications. Stable RPL22-HaloTag, RPL22L1-HaloTag, and HaloTag only expressing HCT116 p53^+/+^ cells were generated using PiggyBac transposon system (System Biosciences) and selected with puromycin (1 µg/ml) for seven days. 1×10^7^ cells in ice-cold PBS to cells were irradiated with 120 mJ/cm^2^ at 254 nm using Stratalinker 2400 (Stratagene). Cells were then harvested, and the pulldown of HaloTag proteins was conducted according to HaloTag Mammalian Pull-Down System protocol (Promega, cat # G6504). Beads were washed thrice with 1× NEB CutSmart Buffer followed by dephosphorylation by using Quick CIP (New England Biolabs, cat # M0525S) and DNase I treatment at 37°C for 30 min. Beads were washed with TRIzol once, thrice with Urea Wash Buffer and thrice with PNK Buffer (New England Biolabs cat # M0201S). 3’ RNA linker ligation and additional denaturing washes using SDS, and urea buffers were conducted according to Gu et al. (Gu et al. 2018) On-bead digestion was performed by adding 100 µL Proteinase K (New England Biolabs, cat # P8107S) followed by incubation at 37°C for 75 min. Resin-bound RNAs were extracted by acid phenol: chloroform: isoamyl alcohol (Invitrogen, cat # AM9720) extraction followed by isopropanol precipitation. RNA pellets were spinned down at 12,000 x *g* for 15 min at 4°C. The pellets were washed twice in 80% ethanol, air dried, and suspended in 8 µL H_2_O. Reverse transcription, gel purification of cDNA and library preparation was as in Gu et al. (Gu et al. 2018) RPL22-HaloTag, RPL22L1-HaloTag, and HaloTag only libraries were sequenced on Illumina NovaSeq X Plus.

### GoldCLIP-seq data analysis

Read 1 for paired-end reads from GoldCLIP libraries containing indexes were first demultiplexed by Ultraplex (v1.2.10) and the adapter sequences were trimmed by Cutadapt (v4.8). The reads were mapped to customized hg38 genomes with a curated rDNA locus (George et al. 2023) using STAR (2.7.10a) with parameters: --genomeLoad NoSharedMemory --readFilesCommand zcat -- outBAMcompression 10 --outFilterMultimapNmax 1 --outFilterMultimapScoreRange 1 -- outFilterScoreMin 10 --outFilterType BySJout --outReadsUnmapped Fastx --outSAMattrRGline ID:foo --outSAMattributes All --outSAMmode Full --outSAMtype BAM Unsorted --outSAMunmapped Within -- runMode alignReads. Mapped reads were deduplicated to remove PCR duplicates using UMI-tools (1.1.5) with parameters: --random-seed 1 --method unique. Mapped reads were split into two bam files with positive or negative strands for peak calling. MACS3 (v3.0.0a6) was used to call peaks with parameters: -g hs -B -keep-dup all --nomodel --extsize 30. Enriched regions were predicted by bdgcmp module using qpois statistic model. Final narrow peak calling was performed using bdgpeakcall function with a cutoff 0.05 for q-value. Mapped reads in peak region were counted using samtools (1.19.2) according to RNA biotypes (lncRNA, miRNA, ncRNA, protein coding, rRNA, snoRNA and snRNA). The genome distribution of binding peaks and metagene plot of peaks at the intron-exon boundary were calculated and visualized by R package GenomicDistributions (1.10.0) (Kupkova et al. 2022). Narrow binding peaks of RPL22 and RPL22L1 were used to generate consensus motifs by findMotifsGenome.pl from HOMER (Hypergeometric Optimization of Motif EnRichment) suite (v4.11) (http://homer.ucsd.edu/homer/) (Heinz et al. 2010). Top five motifs (de novo and known motifs) with reported p-values < 10^-350^ are presented. Integrative Genomics Viewer (v2.17.4) was used to visualize the genomic tracks.

### Enrichment and ontology analyses

Gene set enrichment analysis was performed using GSEA (v4.3.2) (Subramanian et al. 2005). Gene signatures were obtained from the MSigDB ‘‘KEGG_MEDICUS subset of CP’’ collection. Significantly enriched gene sets were ranked and presented using normalized enrichment (NES) score. The gene list from the intersection of RNA-seq and rMATS were subjected to gene enrichment analysis using g:Profiler (https://biit.cs.ut.ee/gprofiler/gost) (Reimand et al. 2007). Top Kyoto Encyclopedia of Genes and Genomes (KEGG) (https://www.genome.jp/kegg/) pathway analyses are reported.

### 45S precursor rRNA chromogenic in situ hybridization (CISH)

Tumors were fixed with 10% formalin, embedded in paraffin and cut to 5 µm. *In situ* hybridization for 5ʹETS precursor rRNA ACD RNAscope 2.5 Brown Kit was conducted as previously described (Guner et al. 2017). Image analysis was conducted on a minimum five tumors/treatment on hematoxylin stained and 5ʹETS CISH assay slides, scanned at x 40 using Ventana DP 200 slide scanner (Roche), and deposited to the Proscia digital pathology database. Necrotic areas were excluded. Image analysis was conducted using HALO v3.0 image analysis platform and Area Quantification module (Indica Labs). The area of brown pixels over the total annotation area (brown+blue) was calculated. The means of these ratios per tumor were used in analyses. All image analyses were conducted in a blinded fashion.

### Mouse tumor models and *in vivo* studies

#### ACUC approval numbers

Mouse studies were conducted according to the animal experimentation permits of the Animal Care and Use Committee at the Johns Hopkins University (MO15M351, MO18M209, MO15M139, MO18M63).

#### Pharmacokinetic and drug uptake analyses

Pharmacokinetic (PK) studies were conducted using a single 30 or 100 mg/kg i.p. or oral doses at Pharmaron Inc. CD-1 mice (n = 3) were sacrificed at various times (5, 15, 45 min, 1.5, 4, 8, 12, 18 and 24 h) and plasma was collected. Plasma samples were analyzed by LC/MS/MS, and PK parameters were determined by non-compartmental analysis of the mean data using WinNonlin version 5.3 (Pharsight Corporation, Mountain View, CA). BOB-42 plasma T_1/2_ was 1.6 hours. For tumor uptake of BOB-42, plasma and tumors were collected 4 hours after the last dose. The samples were analyzed by LC/MS/MS at the Analytical Pharmacology Facility at Sidney Kimmel Comprehensive Cancer Center.

#### A375 human melanoma model

A375 cells (ATCC) were cultured in DMEM supplemented with 10% FBS and 4.5 g/L glucose at 37°C in a humidified atmosphere containing 5% CO_2_. Cells growing in an exponential growth phase were harvested following trypsin-EDTA treatment and counted for tumor inoculation. A375 cells (3 x 10^6^) were suspended in 0.1 mL of PBS and injected subcutaneously at the right upper flank of 6 - 8 weeks old female athymic *nu/nu* mice. Mice bearing subcutaneous A375 tumors (200 mm^3^) were randomized into 4 groups (n = 8 mice/group) using two treatment schemes. Mice were administered i.p. daily for 14 days with vehicle (0.2 M PBS, pH 6.2) or BOB-42 at 12.5 mg/kg, 25 mg/kg or 50 mg/kg. Alternatively, they were administered for 11 days with vehicle (daily, 0.2 M PBS, pH 6.2) or BOB-42 at 50 mg/kg (daily), 75 mg/kg (one day on, one day off) or 100 mg/kg (one day on, two days off). Body weights and tumor sizes were measured thrice per week. Mice were euthanized and blood was collected using cardiopuncture. Blood and plasma were analyzed at the Johns Hopkins Mouse Phenotyping and Pathology Core for clinical chemistry and complete blood count (CBC) using ProCyte Dx Veterinary CBC Hematology Analyzer.

Tumor growth was evaluated by measurement of two perpendicular diameters of tumors with a digital caliper. Tumor volume was expressed in mm^3^ using the formula: V = (L × W2)/2 where L is the tumor length and W is the tumor width. The tumor size was then used to calculate tumor growth inhibition (TGI) using the formula: TGI (%) = [1-(Ti-T0)/ (Vi-V0)] ×100; Ti is the average tumor volume of a treatment group on a given day, T0 is the average tumor volume of the treatment group on the first day of treatment, Vi is the average tumor volume of the vehicle control group on the same day with Ti, and V0 is the average tumor volume of the vehicle group on the first day of treatment.

#### Human colorectal cancer PDX models

PDX models were generated by implanting xenografted human CO-04-0032 (passage 5) and CO-04- 0114 (passage 4) colorectal tumor tissues s.c. into the right flank of 6 - 8 weeks old female *Balb/c* nude mice. Mice with established tumors (150 - 200 mm^3^) were randomized (n = 10 mice/group) and treated i.p. once daily with vehicle (0.2 M PBS, pH 6.2) or BOB-42 (25 mg/kg). Body weights and tumor sizes were measured thrice per week. Mice were euthanized and blood was collected using cardiopuncture for complete blood count (CBC) using Siemens Advia 2120i hematology analyzer. The study was conducted at WuXi AppTec R&D Center, Shanghai, China.

#### Syngeneic mouse colon adenocarcinoma models

MC38 colon tumor cells (cat # NCTT-MC38) purchased from BioVector NTCC Inc. were maintained *in vitro* as a monolayer culture in DMEM medium supplemented with 10% heat inactivated fetal bovine serum, 2 mM glutamine, 100 U/mL penicillin and 100 μg/mL streptomycin at 37°C in an atmosphere of 5% CO2 in the air. Cells growing in an exponential growth phase were harvested following trypsin-EDTA treatment and counted for tumor inoculation. MC38 colon cancer model was established by s.c. inoculation of 0.3 x 10^6^ cells into the right upper flank of 6 - 8 weeks old female *C57BL/6* mice. Treatments were started at day 10 post-tumor inoculation when the average tumor size reached 95 mm^3^. The animals were randomized into 4 groups (n = 10 mice/group) and administered intraperitoneally (i.p.) daily with vehicle (0.2 M PBS, pH 6.2) and BOB-42 25 mg/kg, or biweekly with mouse anti-PD-1 antibody RMP1-14 (BioXcell, cat # BE0146) 10 mg/kg, or using a combination of BOB-42 and mouse anti-PD-1 antibody as above.

Body weights and tumor sizes were measured thrice per week throughout the experiment. All animals were monitored at least once daily. Survival endpoints were reached and recorded when the tumor volume reached 3,000 mm^3^, or mouse health concern or loss of 20% body weight, whichever occurred first. Mice were euthanized by standard CO_2_ asphyxiation. Kaplan-Meier and log-rank test were used to compare survival curves between groups. All data were analyzed using GraphPad Prism software 8.0.2. *P* < 0.05 was considered statistically significant. The study was conducted at WuXi AppTec R&D Center, Shanghai, China. All animal studies including the mouse euthanasia procedure were performed according to the guidelines approved by the Institutional Animal Care and Use Committee (IACUC) of WuXi AppTec following the guidance of the Association for Assessment and Accreditation of Laboratory Animal Care (AAALAC).

#### Tumor infiltrating leukocyte (TIL) assessment by flow cytometry

At day 6 post-implantation of MC38 colon cancer cells, tumor-harboring mice (average tumor volume 49 mm^3^) were randomized into 4 groups (n = 7 mice/group). Mice were administered (i.p.) with vehicle (0.2 M PBS, pH 6.2), BOB-42 25 mg/kg (dissolved in 0.2 M PBS, pH 6.2), mouse anti-PD-1 antibody RMP1-14, 10 mg/kg (diluted in 0.0067M PBS, pH 7.4) (BioXcell, cat # BE0146), or the combination of BOB-42 and mouse anti-PD-1 antibody. Tumors were excised 24 h after the last dose. Tumors were minced, dissociated into single-cell suspension and filtered through 70 μm cell strainers. Cells were washed twice with DPBS, then resuspended in PBS, and seeded into 96-well plates (1×10^6^ cells per well). Brilliant Violet 421 live/dead dye (0.1 µL per well) was added and incubated for 30 min at 4°C in the dark. Cells were washed and suspended with 100 µL staining buffer. Purified rat anti-mouse CD16/CD32 antibody (BD Biosciences, cat # 553142) was added and incubated at 4°C for 5 min. Stained cells were then incubated with fluorochrome Brilliant UltraViolet 395-conjugated CD8 (BD Biosciences, cat # 565968), Brilliant Violet 605-conjugated anti-mouse CD19 (BD Biosciences, cat # 563148), Brilliant Violet 711 conjugated anti-mouse CD11b (Biolegend, cat # 101242), FITC-conjugated anti-mouse CD335 (Biolegend, cat # 137606), PerCP-conjugated anti-mouse CD4 (Biolegend, cat # 100432), PE-conjugated anti-mouse PD-L1 (Biolegend cat # 124307), APC-conjugated anti-mouse PD-1 (Biolegend, cat # 135209), Alexa Fluor 700-conjugated anti-mouse CD45 (BD Biosciences, cat # 560510), and APC-Cy7-conjugated anti-mouse CD3 (Biolegend, cat # 100221) antibodies for 30 min at 4°C. After staining of the extracellular antigens, cells were washed twice with staining buffer, fixed and permeabilized for 30 minutes and stained with PE-Cy7-conjugated monoclonal Foxp3 antibody (eBioscience, cat # 25-5773-82). Stained cells were washed twice again and suspended in 200 µL staining buffer. Flow cytometry was conducted using BD FACS LSR Fortessa X20 Flow Cytometer and FlowJo software.

#### Statistical analyses and reproducibility

No data were excluded from the analyses. Data distribution was assumed normal, but this was not formally tested. Mouse treatment groups were randomized throughout the study. There was no randomization of the *in vitro* studies. Data collection and performance were unblinded, unless otherwise stated.

## SUPPLEMENTAL INFORMATION

**Document S1.** Supplemental Figures S1 – S6 and Tables S1, S2, S3 and S4.

**Document S2.** Uncut immunoblotting and Tapestation RNA gels.

**Supplemental Tables S5-S11 each are provided as separate files due to their size:**

Table S5. Splicing events in RPL22^RE^ cells. Data related to Figure 5 and Figure S4.

Table S6. Splicing events in RPL22L1^KO^ cells. Data related to Figure 5 and Figure S4.

Table S7. BMH-21 induced splicing events in HCT116 p53^+/+^ cells. Data related to Figure 6A.

Table S8. BMH-21 induced splicing events in HCT116 p53^-/-^ cells. Data related to Figure 6B.

Table S9. BMH-21 induced splicing events in A375 cells. Data related to Figure 6C.

Table S10. Peak RNA-binding sites of RPL22^Halo^. Data related to Figure 7 and Figure S6.

Table S11. Peak RNA-binding sites of RPL22L1^Halo^. Data related to Figure and Figure S6.

